# Top-down control of hippocampal signal-to-noise by prefrontal long-range inhibition

**DOI:** 10.1101/2021.03.01.433441

**Authors:** Ruchi Malik, Yi Li, Selin Schamiloglu, Vikaas S. Sohal

## Abstract

The prefrontal cortex (PFC) is postulated to exert ‘top-down control’ by modulating information processing throughout the brain to promote specific actions based on current goals. However, the pathways mediating top-down control remain poorly understood. In particular, knowledge about direct prefrontal connections that might facilitate top-down prefrontal control of information processing in the hippocampus remains sparse. Here we describe novel monosynaptic long-range GABAergic projections from PFC to hippocampus. These preferentially inhibit vasoactive intestinal polypeptide expressing interneurons, which are known to disinhibit hippocampal microcircuits. Indeed, stimulating prefrontal–hippocampal GABAergic projections increases hippocampal feedforward inhibition and reduces hippocampal activity *in vivo*. The net effect of these actions is to specifically enhance the signal-to-noise ratio for hippocampal representations of objects. Correspondingly, stimulation of PFC-to-hippocampus GABAergic projections promotes object exploration. Together, these results elucidate a novel top-down pathway in which long-range GABAergic projections target disinhibitory microcircuits, thereby enhancing signals and network dynamics underlying exploratory behavior.

## Introduction

The prefrontal cortex (PFC) plays a crucial role in executive functions and the top-down control of brain activity and behavior (Gazzaley and D’Esposito, 2007; Miller and Cohen, 2001; Miller, 2000). It is postulated that PFC bidirectionally communicates with several cortical and subcortical brain regions, monitoring and gating their ongoing activity, in order to exert top-down executive control over behavior. One brain region that is known to bidirectionally interact with the PFC is the hippocampus (HPC), a key brain region for processing spatial information, and using spatial representations to guide behavior. Accumulating evidence in humans and animal models highlights an essential role of network interactions between the PFC and HPC in cognitive and emotional behaviors (Eichenbaum, 2017; Jin and Maren, 2015; Preston and Eichenbaum, 2013; Shin and Jadhav, 2016; Sigurdsson and Duvarci, 2016; Yu and Frank, 2015). Importantly, abnormal PFC-HPC interactions are thought to contribute to cognitive and emotional deficits in several neuropsychiatric disorders, including schizophrenia, depression, and anxiety disorders (Cunniff et al., 2020; Godsil et al., 2013; Kupferschmidt and Gordon, 2018; Li et al., 2015; Sigurdsson et al., 2010). Owing to the importance of PFC-HPC interactions in normal and pathological behaviors, much work has been focused on elucidating how these regions interact.

Functional imaging studies in humans as well as rodent studies using lesions and pharmacological inactivation have shown that concurrent activity in, and communication between, the PFC and HPC is essential for spatial exploratory behaviors (Bähner et al., 2015; Churchwell et al., 2010; DeVito and Eichenbaum, 2010; Floresco et al., 1997; Wang and Cai, 2006; Yoon et al., 2008). Neural activity and network oscillations synchronize across the PFC and HPC during spatial exploratory behaviors (Colgin, 2011; Jones and Wilson, 2005; O’Neill et al., 2013; Spellman et al., 2015). In particular, oscillatory activity in HPC leads PFC activity when rats explore spatial contexts, but this pattern of synchronization switches to PFC leading when rats explore objects in their environment (Place et al., 2016) or arrive at decision points in a maze (Hallock et al., 2016). Furthermore, inactivating the PFC alters the encoding of spatial information in the HPC (Guise and Shapiro, 2017; Kyd and Bilkey, 2003). These findings suggest that the PFC exerts top-down control over HPC activity at key behavioral timepoints, but knowledge about direct anatomical projections that mediate this kind of top-down prefrontal control is lacking. In fact, whereas much is known about the direct anatomical pathways from the HPC-to-PFC (Hoover and Vertes, 2007; Jay and Witter, 1991), most top-down communication in the PFC-to-HPC direction is thought to occur indirectly, via the thalamic nucleus reuniens (NR) (Hoover and Vertes, 2012; Vertes et al., 2007; Xu and Südhof, 2013). Not only are the anatomical substrates for top down control unknown, the manner in which top down control operates is also unclear. I.e., does the PFC exerts top-down control by transmitting specific information, e.g., representations of specific actions or goals, or alternatively, does it modulate the network state, changing the nature of emergent circuit computations in downstream regions?

Previous studies of PFC-HPC interactions have focused on direct and indirect excitatory (glutamatergic) connections between these structures. Growing evidence indicates that cortical circuits also include specialized populations of long-range projecting GABAergic (LRG) inhibitory neurons (Jinno et al., 2007; Melzer and Monyer, 2020; Tamamaki and Tomioka, 2010). In some cases, these LRG projections have been shown to control oscillatory synchronization between structures (Christenson Wick et al., 2019; Francavilla et al., 2018; Melzer et al., 2012), suggesting that they may be important regulators of interregional communication. We recently reported that the PFC also contains specialized LRG projection neurons capable of influencing behavior (Lee et al., 2014). Therefore, we hypothesized that a specialized population of PFC LRG projection neurons might serve as the anatomical substrate through which the PFC exerts top-down control over hippocampal information processing.

Here, we report a novel population of LRG neurons in the PFC that send direct inhibitory projections to the dorsal hippocampus (dHPC). Notably, these prefrontal LRG projections target local disinhibitory microcircuits and modulate network oscillations in dHPC. Through these actions, PFC–dHPC LRG projections promote network states associated with object exploration, enhance hippocampal representations of object locations, and elicit corresponding increases in the time mice spend exploring objects. Together, our results show how the PFC exerts top-down control over information processing in the HPC by acting through a novel circuit motif: long-range GABAergic projections which inhibit disinhibitory microcircuits, thereby altering emergent network dynamics and promoting specific exploratory behaviors.

## Results

### Hippocampus projecting long-range GABAergic (LRG) neurons in the PFC

To label potential PFC-to-HPC LRG projections, we used *Dlxi12b-Cre* mice, which specifically express Cre recombinase in GABAergic neurons (Lee et al., 2014; Potter et al., 2009). We injected an adeno-associated virus (AAV) to drive Cre-dependent expression of the fluorescent reporter eYFP (AAV5-EF1α-DIO-eYFP) in the PFC of *Dlxi12b-Cre* mice (Fig. 1A). After waiting 6–8 weeks for viral transduction, we observed robust eYFP expression in the cell bodies of GABAergic neurons in the PFC and also observed many axonal fibers in the CA1 and dentate gyrus subfields of dHPC (Fig. 1B and Fig. S1). Importantly, no eYFP+ cell bodies were observed in the HPC.

**Figure 1:**
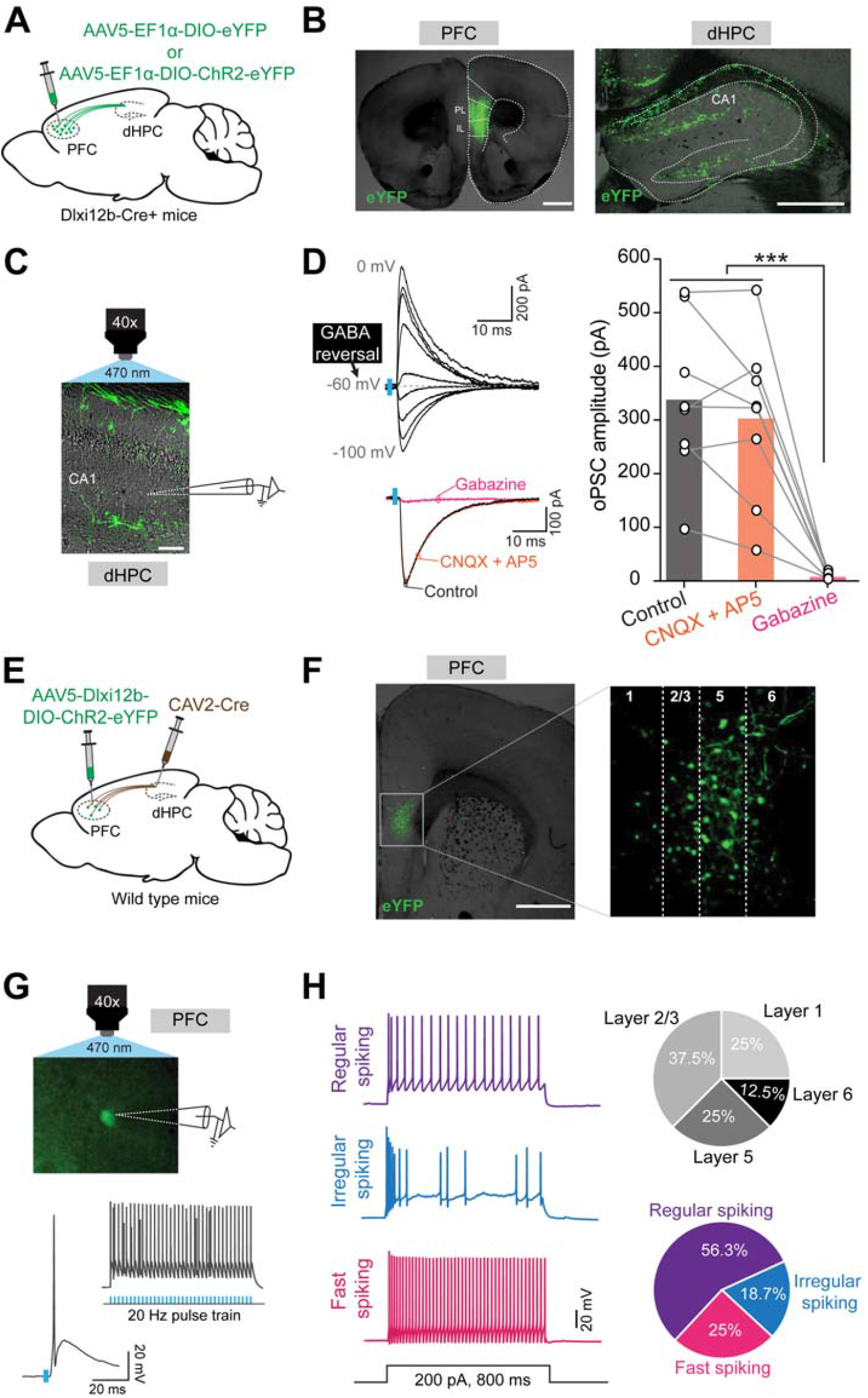
A heterogeneous population of PFC inhibitory neurons sends direct LRG projections to the dHPC. (**A)** Schematic illustrating the anterograde tracing strategy. Cre-dependent eYFP or ChR2-eYFP virus was injected into the PFC of *Dlxi12b-Cre*+ mice. **(B)** Representative images showing eYFP+ PFC GABAergic neurons (left) and eYFP+ axonal fibers in the dHPC (right). Scale bars, 1 mm and 0.5 mm, respectively. Prelimbic cortex (PL), infralimbic cortex (IL), and hippocampal CA1 regions are labeled. **(C)** Overlaid fluorescent and DIC images of a hippocampal section showing ChR2-eYFP+ axonal fibers (green) in dorsal CA1. During *ex-vivo* patch clamp recordings from hippocampal neurons, ChR2+ LRG axons were optogenetically activated by pulses of blue light (5 ms, 470 nm) delivered through the 40x objective. Scale bar, 100 µm. **(D)** Top left: example traces showing that optogenetically evoked postsynaptic currents (oPSCs) in recipient CA1 neurons reverse at the GABA reversal potential (gray dashed line). Blue bars denote light pulses. Bottom left: example oPSCs recorded from a CA1 neuron in control aCSF (black), after adding CNQX + AP5 (orange), and after adding Gabazine (magenta). Right: oPSC amplitudes were significantly reduced by Gabazine. Open circles represent data from individual neurons (n = 8) and bars represent averages; one-way ANOVA followed by Tukey’s multiple comparison test, *** p < 0.001. **(E)** Schematic demonstrating the intersectional strategy to target dHPC-projecting PFC LRG neurons. Retrogradely transducing canine adenovirus type-2 Cre (CAV2-Cre) was injected into dHPC, and a Cre-dependent AAV that drives expression of ChR2-eYFP using a GABAergic neuron-specific enhancer (Dlxi12b) was injected into PFC. **(F)** Representative images showing ChR2-eYFP expression in dHPC projecting GABAergic neurons in PFC. White dotted box in the left image corresponds to the magnified image shown in right. Numbers indicate the cortical layers. Scale bar, 1 mm. **(G)** Top: representative image showing *ex-vivo* patch clamp recording obtained from ChR2-eYFP expressing PFC–dHPC LRG neuron. Bottom: example traces showing PFC–dHPC LRG neuron firing elicited in response to a single light pulse (5 ms, 470 nm) or a 20 Hz train. **(H)** Left: example voltage responses of PFC–dHPC LRG neurons to depolarizing current injections. Top right: pie chart showing the laminar distribution of recorded PFC–dHPC LRG neurons. Bottom right: pie chart showing the percentage of PFC–dHPC LRG neurons with various physiological properties. See also **Figures S1, S2** and **Table S1**.

Next, we asked whether these PFC LRG axon terminals synapse onto neurons in the dHPC. To address this, we injected AAV into the PFC of *Dlxi12b-Cre* mice to drive Cre-dependent expression of Channelrhodopsin-eYFP (ChR2-eYFP) in PFC GABAergic neurons, then, after waiting for expression, made recordings from acute hippocampal slices (Fig. 1C). Notably, optogenetic activation of PFC LRG axonal fibers in dHPC slices elicited robust short-latency postsynaptic currents (oPSCs) in dHPC neurons. These currents reversed at the GABA reversal potential, were not affected by glutamatergic receptor antagonists, and were completely blocked by bath application of the GABA_A_ receptor antagonist gabazine (10 µM) (Fig. 1D).

Following the identification of PFC–dHPC LRG projections, we asked whether these dHPC-projecting PFC LRG neurons have distinct electrophysiological and molecular properties, and whether these neurons are located in superficial or deeper cortical layers of the PFC. To address these questions, we used an intersectional strategy to selectively express ChR2-eYFP in dHPC-projecting PFC LRG neurons. Specifically, we injected two viruses: a retrogradely transducing canine adenovirus type-2 Cre (CAV2-Cre) into dHPC, and a Cre-dependent AAV expressing ChR2-eYFP under control of the *Dlxi12b* enhancer into PFC (Lee et al., 2014) (Fig. 1E, F). We then made *ex-vivo* patch clamp recordings from dHPC-projecting LRG neurons in PFC (identified by eYFP expression), and recorded reliable short-latency light-evoked action potentials (APs) to confirm that they were ChR2-expressing (Fig. 1G). These recordings revealed that the dHPC-projecting PFC LRG neuronal population is electrophysiologically diverse, comprising neurons with regular spiking (9/16 neurons), irregular spiking (3/16 neurons), and fast spiking (4/16 neurons) physiological properties (Fig. 1H and Table S1). dHPC-projecting PFC LRG neurons were distributed across superficial and deeper layers of the prelimbic (PL) portion of the PFC. By combining injection of a retrograde tracer (Alexa 594-tagged cholera toxin, CTb) in dHPC with immunohistochemistry in PFC (Fig. S2A, B), we found that dHPC-projecting PFC LRG neurons include parvalbumin (PV), somatostatin (SST), and vasoactive intestinal polypeptide (VIP)-expressing subpopulations (Fig. S2C, D). We also observed small percentages of calretinin (CR) and neuropeptide-Y (NPY) expressing PFC–dHPC LRG neurons. However, none of the PFC LRG neurons in our study showed immunoreactivity for neuronal nitric oxide synthase (nNOS). Taken together, these results reveal that the dHPC receives direct LRG projections which originate from a heterogeneous population of GABAergic inhibitory neurons located across multiple layers of the PFC.

### PFC LRG projections target hippocampal disinhibitory interneurons

Next, we asked how electrophysiologically and molecularly heterogeneous PFC LRG projection neurons affect circuit computations in the CA1 subregion, which is the primary output region of the dHPC. Specifically, we asked whether PFC LRG projections target specific cell-types in the CA1 subregion. We expressed ChR2-eYFP in PFC LRG projections and obtained *ex-vivo* patch clamp recordings from excitatory pyramidal neurons and GABAergic interneurons located in different topographical layers of CA1 subregion in acute hippocampal slices (Fig. 2A). Interestingly, we observed robust optogenetically-evoked IPSCs in CA1 interneurons (55/70 connected, henceforth referred to as recipient interneurons), but not in CA1 pyramidal neurons (PNs; 0/38 connected) (details of interneuron and PN classification in Methods). Notably, many of the recipient CA1 interneurons were located near the border between stratum radiatum (SR) and stratum lacunosum-moleculare (SLM) (Fig. 2B). Furthermore, recipient CA1 interneurons comprised physiologically heterogeneous subtypes including regular spiking, irregular spiking, and fast spiking interneurons (Fig. 2B and Table S2). In order to determine whether the PFC LRG projections target molecularly defined interneuron subtypes in CA1, we filled a subset of the recipient interneurons with biocytin and quantified the immunoreactivity for three molecular markers commonly expressed in CA1 interneurons– PV, SST, and VIP. Surprisingly, we found that a majority of recipient interneurons expressed VIP (7/11). By contrast, none of the recipient interneurons we examined expressed PV or SST (0/10) (Fig. 2C).

**Figure 2:**
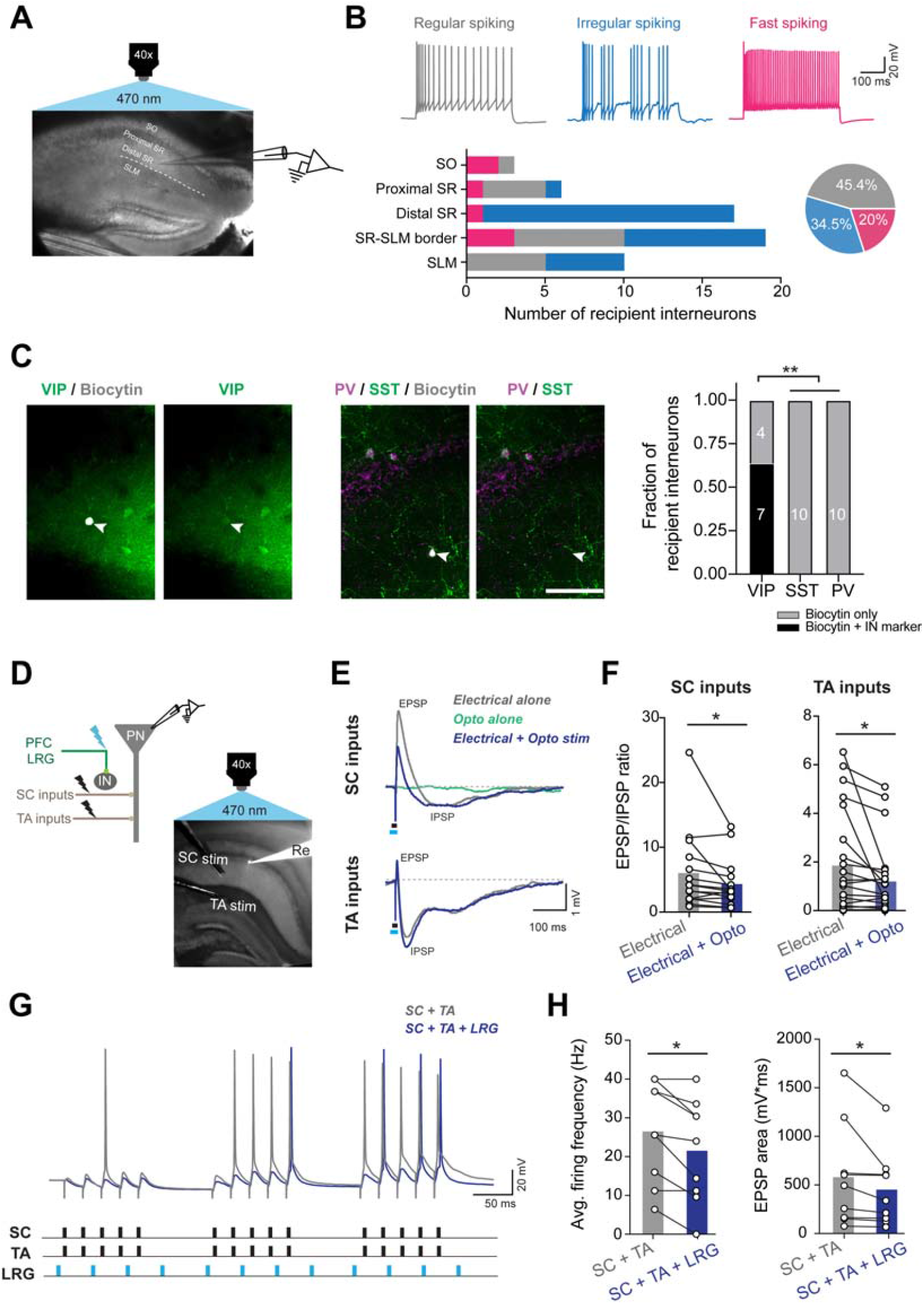
PFC LRG projections preferentially target interneuron-selective interneurons (ISIs) and increase feedforward inhibition in the CA1 microcircuit. **(A)** Example DIC image of a hippocampal slice showing *ex-vivo* patch clamp recording from a CA1 neuron during optogenetic stimulation of PFC LRG projections. Different layers of CA1 are labeled; stratum oriens (SO), stratum radiatum (SR), and stratum lacunosum-pyramidale (SLM); dashed white line represents the border between SR and SLM. **(B)** Top: example voltage responses to depolarizing current injections for CA1 neurons which receive input from PFC LRG projections. Regular spiking (gray), irregular spiking (blue), and fast spiking properties (magenta) are observed among recipient CA1 neurons. Bottom left: number of recipient interneurons with fast spiking, regular spiking and irregular spiking physiology at different laminar locations in CA1 is plotted. Bottom right: pie chart showing the percentage of recipient CA1 neurons with regular spiking (25/55), irregular spiking (19/55), and fast spiking (11/55) properties. **(C)** Left: Representative images showing staining for inhibitory neuron (IN) markers (VIP, PV or SST) in recipient CA1 neurons filled with biocytin. Scale bar, 100 µm. Right: fraction of recipient neurons which stained positive for VIP or (in separate sections) for PV or SST. More recipient neurons stained positive for VIP vs. PV or SST (Chi-square test, ** p < 0.01). **(D)** Top left: Schematic showing the experimental configuration. Right: example hippocampal image showing Alexa-594 filled recording electrode (Re) targeting a CA1 pyramidal neuron (PN). Stimulating electrodes were placed in SR and SLM to stimulate Schaffer collateral (SC stim) or temporoammonic (TA stim) inputs, respectively. Brief pulses of blue light (5ms, 470 nm) delivered through the 40x objective were used to optogenetically stimulate PFC LRG projections. **(E)** Excitatory and inhibitory postsynaptic potentials (EPSPs and IPSPs) elicited by electrical stimulation of SC or TA inputs (black bar) in the presence or absence of optogenetic stimulation of PFC–dHPC LRG projections (cyan bar). Gray traces show responses to electrical stimulation alone, blue traces show responses during combined electrical + optogenetic stimulation, and green trace shows response to optogenetic stimulation alone. **(F)** Right: optogenetic stimulation of PFC–dHPC LRG projections significantly decreased EPSP to IPSP ratios for both SC (n = 15 cells) and TA (n = 22 cells) inputs. Lines connect values from individual neurons and bars represent averages; two-way paired t-test, ** p < 0.01, * p < 0.05. **(G)** Example voltage responses of CA1 PN to coincident theta-burst stimulation (TBS) of SC and TA inputs (black bars) combined with 20 Hz optogenetic stimulation of PFC LRG projections (cyan bars). Gray trace shows voltage response to SC and TA electrical stimulation, and blue trace denotes response during SC and TA electrical stimulation + optogenetic stimulation of PFC LRG projections. **(H)** Average firing frequency (left) and EPSP area (right) during TBS of SC and TA inputs are reduced by concomitant optogenetic stimulation of PFC LRG projections. Open circles represent individual neurons (n = 9) and bars represent averages; two-way paired t-test, * p < 0.05. See also **Figure S3** and **Table S2**.

### PFC LRG projections regulate excitatory input integration in CA1 microcircuit

Since VIP is predominantly expressed by interneuron-selective interneurons (ISIs) which produce circuit disinhibition in CA1 (Acsády et al., 1996a; 1996b; Chamberland and Topolnik, 2012; Turi et al., 2019), we hypothesized that PFC–dHPC LRG projections may inhibit VIP+ ISIs, thereby reducing disinhibition and increasing feedforward inhibition in the CA1 microcircuit. To test this prediction, we quantified the effect of optogenetic stimulation of PFC LRG projections on excitatory and inhibitory postsynaptic potentials (EPSPs and IPSPs) elicited by two major afferent input pathways: Schaffer collateral (SC) and temporoammonic (TA) inputs (Fig. 2D). Specifically, during *ex-vivo* patch clamp recordings from CA1 PNs, we delivered electrical stimulation to SC or TA inputs concomitant with optogenetic stimulation of ChR2+ PFC–dHPC LRG axon fibers. While the optogenetic stimulation of PFC–dHPC LRG axons alone did not elicit discernable postsynaptic potentials in CA1 PNs, concomitant electrical and optogenetic stimulation significantly increased the size of IPSPs relative to EPSPs for both SC and TA inputs (Fig. 2E, F and Fig. S3A,B). This reduction in the excitation to inhibition ratio (E/I ratio) for SC and TA inputs is consistent with our prediction and suggests that the PFC LRG projections increase feedforward inhibition by inhibiting disinhibitory VIP+ interneurons in CA1.

We then asked whether increased feedforward inhibition during stimulation of PFC LRG projections affects the input-output transformation performed by CA1 PNs. Coincident activation of SC and TA input pathways, often in a theta-burst stimulation (TBS) pattern, is known to cause supralinear input summation and spiking in CA1 PNs (Ang et al., 2005; Bittner et al., 2015; Malik and Johnston, 2017). This nonlinear input integration and coincidence detection in CA1 PNs is tightly regulated by the activity of CA1 interneurons and considered crucial for hippocampal information processing (Grienberger et al., 2017; Milstein et al., 2015). To determine how PFC–dHPC LRG projections modulate input integration in CA1 PNs, we combined electrical TBS of SC and TA inputs with optogenetic stimulation of PFC–dHPC LRG projections (20 Hz, 5 ms pulses) (Fig. 2G). Again, consistent with increased feedforward inhibition, optogenetic stimulation of LRG projections reduced firing and EPSP summation during TBS (Fig. 2H). Importantly, firing of CA1 PNs in response to depolarizing current injections (i.e., neuronal depolarization without recruitment of microcircuit inhibition) was not affected by optogenetic stimulation of these PFC LRG projections (Fig. S3C). Taken together, our *ex-vivo* electrophysiological analyses show how PFC–dHPC LRG projections regulate synaptic integration and input-output gain by enhancing feedforward inhibition onto CA1 PNs (Fig. S3D).

### PFC–dHPC LRG projections promote object exploration

Communication between PFC and HPC is implicated in many spatial and object exploration behaviors (DeVito and Eichenbaum, 2010; Jin and Maren, 2015; Preston and Eichenbaum, 2013; Spellman et al., 2015; Yu and Frank, 2015). Notably, both structures synchronize at theta frequency with dHPC leading when rodents enter a spatial context, but the directionality switches to PFC leading when animals sample an object (Place et al., 2016). This suggests an important role for top-down communication from PFC to dHPC during object exploration. Therefore, we quantified how PFC–dHPC LRG projections affect object exploration in freely behaving mice. Optogenetic stimulation of PFC–dHPC LRG projections (20 Hz, 5 ms pulses, 473 nm, ∼3-4 mW) dramatically increased the time *Dlxi12b-Cre* mice spent engaged in novel object exploration (NOE) (Fig. 3A, B). Light delivery alone had no effect in control (Cre-negative) mice. Increases in NOE occurred during both early and late portions of the testing session (Fig. 3C, D), and reflected increased numbers of both short- and long-duration bouts of object exploration (Fig. 3E). Optogenetic stimulation of PFC–dHPC LRG projections did not affect the distance travelled in an open field, time spent on the stimulated side during a real-time place preference task, or the time spent exploring a novel juvenile mouse (Fig. S4A–C). Thus, activating PFC–dHPC LRG projections specifically increases NOE without nonspecifically affecting movement or other exploratory behaviors.

**Figure 3:**
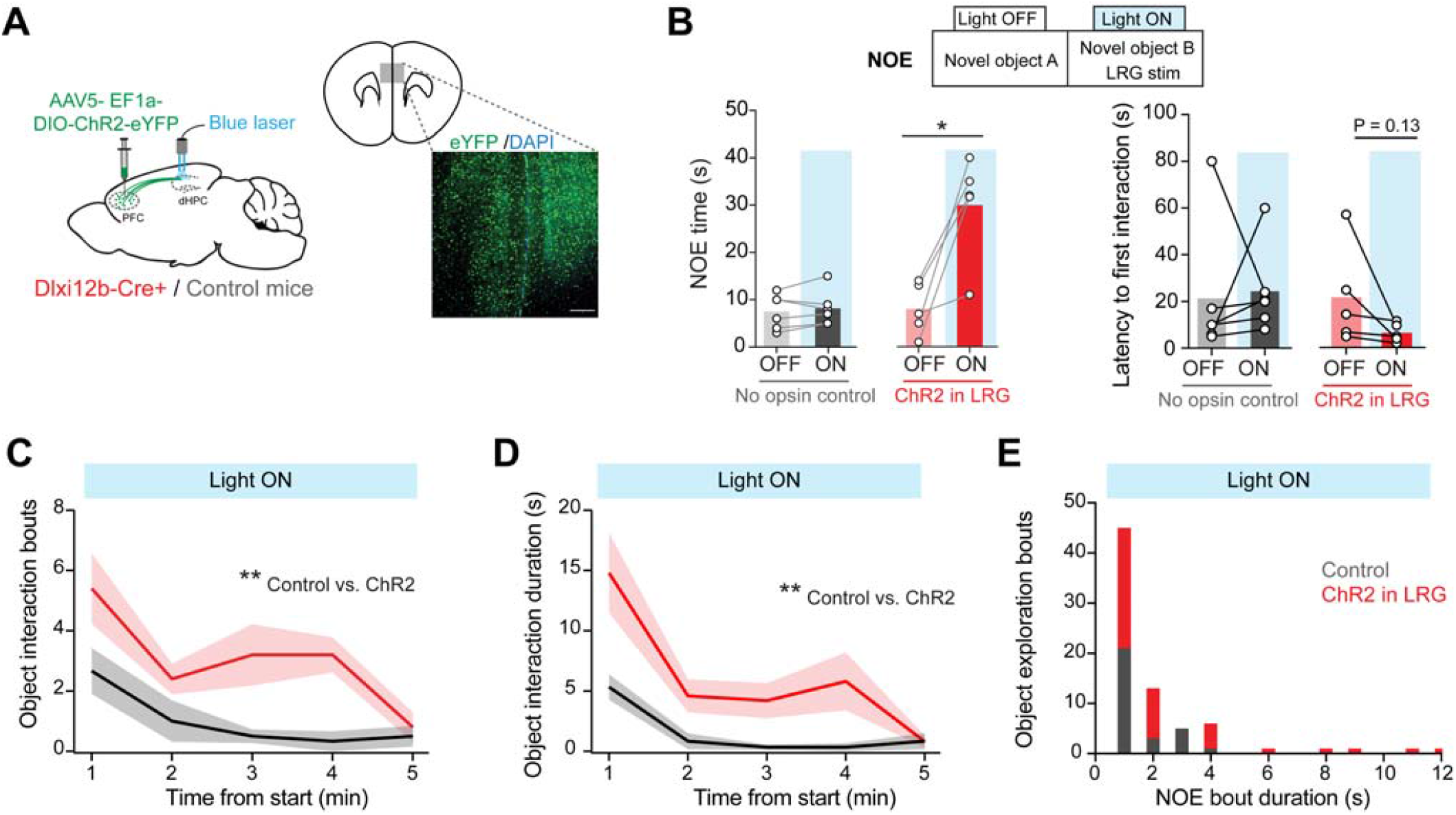
Activating PFC–dHPC LRG projections increases novel object exploration. **(A)** Left: schematic illustrating the experimental design. Bilateral injections of Cre-dependent ChR2-eYFP virus into the PFC of *Dlxi12b-Cre*+ mice or *Cre-negative* (control) mice. Bilateral optical fibers were implanted over dHPC. Right: representative image showing ChR2-eYFP expression in PFC GABAergic neurons. Scale bar, 200µm. **(B)** Top: novel object exploration (NOE) was measured in the presence (Light ON) or absence (Light OFF) of optogenetic stimulation (473nm, 1.5–2 mW/fiber, 5 ms pulses at 20 Hz). Bottom left: optogenetic stimulation significantly increased NOE time in ChR2+ mice (n = 5) but not in control mice (n = 6). Bottom right: Latency to first interaction with a novel object for ChR2+ and control mice is plotted. Open circles represent values from individual mice and bars indicate averages. Two-way paired t-test, * p < 0.05. **(C–D)** Number of object interaction bouts within 1-minute bins (**C**), and duration of object interaction within 1-minute bins (**D**), over the duration of a 5-minute NOE testing session. Two-way repeated measures ANOVA, ** p < 0.01. **(E)** Distribution of NOE bout durations during optogenetic stimulation of PFC LRG projections in control and ChR2-expressing mice. See also **Figure S4**.

### PFC LRG projections promote network oscillations associated with object exploration

Next, we explored potential circuit mechanisms through which PFC–dHPC LRG projections might impact NOE. We did this in two ways. First, we recorded local field potentials (LFPs) to determine whether stimulation of PFC–dHPC LRG projections might induce network states conducive to NOE (Fig. 4A). In comparison to baseline home cage (HC) exploration, NOE recruited synchronized oscillations in the low-gamma (25–55 Hz) band across the PFC-dHPC network. Specifically, during NOE we observed a significant increase in low-gamma power in both structures as well as an increase in low-gamma phase synchrony between the PFC and dHPC (Fig. 4B). While the increase in low-gamma activity was most prominent, NOE was also associated with significant increases in power (but not synchrony) for high-gamma activity (both structures) and theta activity (dHPC only) (Fig. S5A, B). The NOE related change in low-gamma frequency oscillations is particularly notable because previous studies have shown that object exploration increases low-gamma synchrony between hippocampal subfields (Trimper et al., 2017). Since microcircuit interactions between local CA1 interneurons and PNs are known to critically regulate gamma oscillations (Csicsvari et al., 2003; Tukker et al., 2007), we hypothesized that by modulating microcircuit inhibition, PFC–dHPC LRG projections could contribute to NOE-associated changes in gamma activity. To test whether PFC–dHPC LRG projections might support these changes in network activity, we combined optogenetic stimulation with multisite LFP recordings in *Dlxi12b-Cre* mice expressing ChR2 in PFC–dHPC LRG projections (Fig. 4C). Indeed, optogenetic stimulation of PFC LRG terminals (20 Hz, 5 ms pulses, 473 nm, ∼3-4 mW) in dHPC mimicked the increases in both low-gamma LFP power and low-gamma phase synchrony observed during NOE (Fig. 4D and Fig. S5C, D). Thus, PFC LRG projections promote a network state associated with object exploration, a behavior known to rely on top-down PFC–dHPC communication.

**Figure 4:**
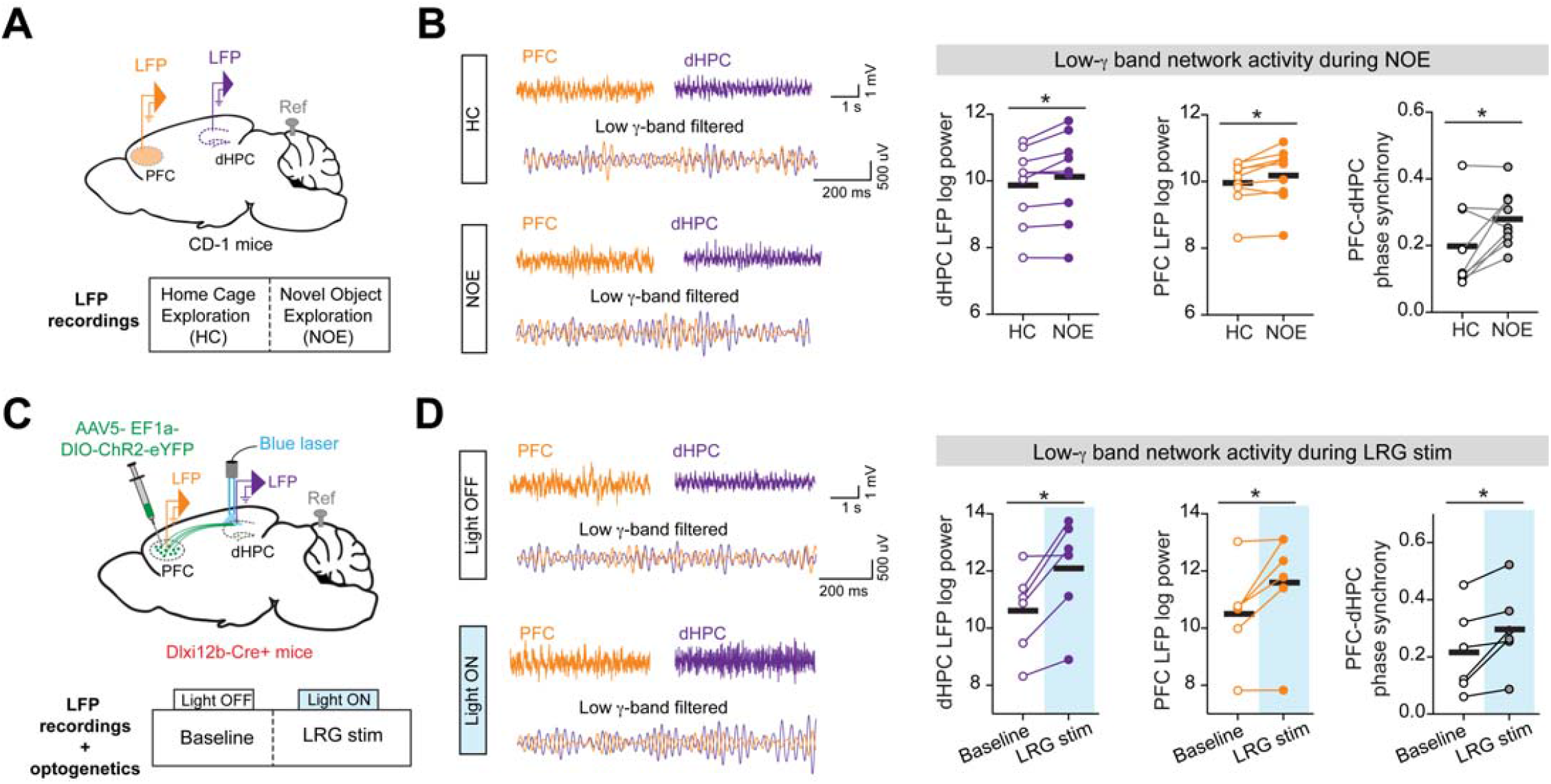
Activating PFC–dHPC LRG projections increases low-gamma oscillations associated with NOE. **(A)** Top: LFP electrodes were implanted in PFC and dHPC, reference electrode was implanted over the cerebellum. Bottom: LFPs recorded while mice were in their home cages (HC) were compared to LFPs recorded during NOE epochs. **(B)** Left: example raw and low-gamma (low-γ) frequency filtered LFPs recorded during HC and NOE epochs are shown. Right: low-γ power in dHPC and PFC and low-γ phase synchrony between PFC and dHPC were all significantly higher during NOE. Open and filled circles represent data from individual mice (n = 9) and solid black lines represent averages. Two-way paired t-test, * p < 0.05. **(C)** Top: Schematic illustrating the experimental design for combined optogenetic stimulation and LFP recordings. Cre-dependent ChR2-eYFP virus was bilaterally injected into the PFC of *Dlxi12b-Cre*+ mice. Bilateral optical fibers were implanted over dHPC; LFP electrodes were implanted in PFC and dHPC; reference electrode was implanted over the cerebellum. Bottom: LFPs were recorded during baseline epochs (Light OFF) or LRG stimulation epoch (Light ON; 473 nm, 5 ms pulses at 20 Hz). **(D)** Left: example raw and low-γ band filtered LFPs recorded during Light OFF and Light ON epochs. Right: low-γ power in dHPC and PFC and low-γ phase synchrony between PFC and dHPC were all significantly higher during LRG stim (Light ON) epoch. Open and filled circles represent data from individual mice (n = 6) and solid black lines represent averages. Two-way paired t-test, * p < 0.05. See also **Figure S5**.

### PFC–dHPC LRG projections reduce hippocampal activity *in vivo*

Having established that PFC–dHPC LRG projections promote network states associated with NOE as well as NOE itself, we next studied how these projections affect NOE-associated hippocampal activity at the level of single cells. For this, we expressed jGCaMP7f in dHPC CA1 neurons (Dana et al., 2019) and used one-photon miniaturized microscopes (miniscopes) to record *in vivo* neuronal Ca^2+^ activity while mice explored novel objects. Concurrently, we expressed the red-shifted excitatory opsin ChrimsonR (Klapoetke et al., 2014; Stamatakis et al., 2018) in PFC–dHPC LRG projections (Fig. 5A–C and Fig. S6A, B). On day 1, mice explored a novel object in the absence of optogenetic stimulation of PFC LRG projections. Across all neurons, Ca^2+^ activity decreased significantly during NOE, relative to the HC epoch (Fig. 5D), although a small fraction of neurons (13/55 neurons) had higher activity during NOE (Fig. S6C). On day 2, we optogenetically stimulated PFC–dHPC LRG projections during both HC and NOE epochs. Stimulating LRG projections during the HC epoch significantly reduced activity (compared to the pre-stimulation HC period). Activity was then further reduced when the mice subsequently engaged in NOE (Fig. 5D and Fig. S6C). This overall reduction in population activity *in vivo* is consistent with our *ex-vivo* observation that activating PFC–dHPC LRG projections tends to enhance feedforward inhibition and reduce spiking in CA1 PNs.

**Figure 5:**
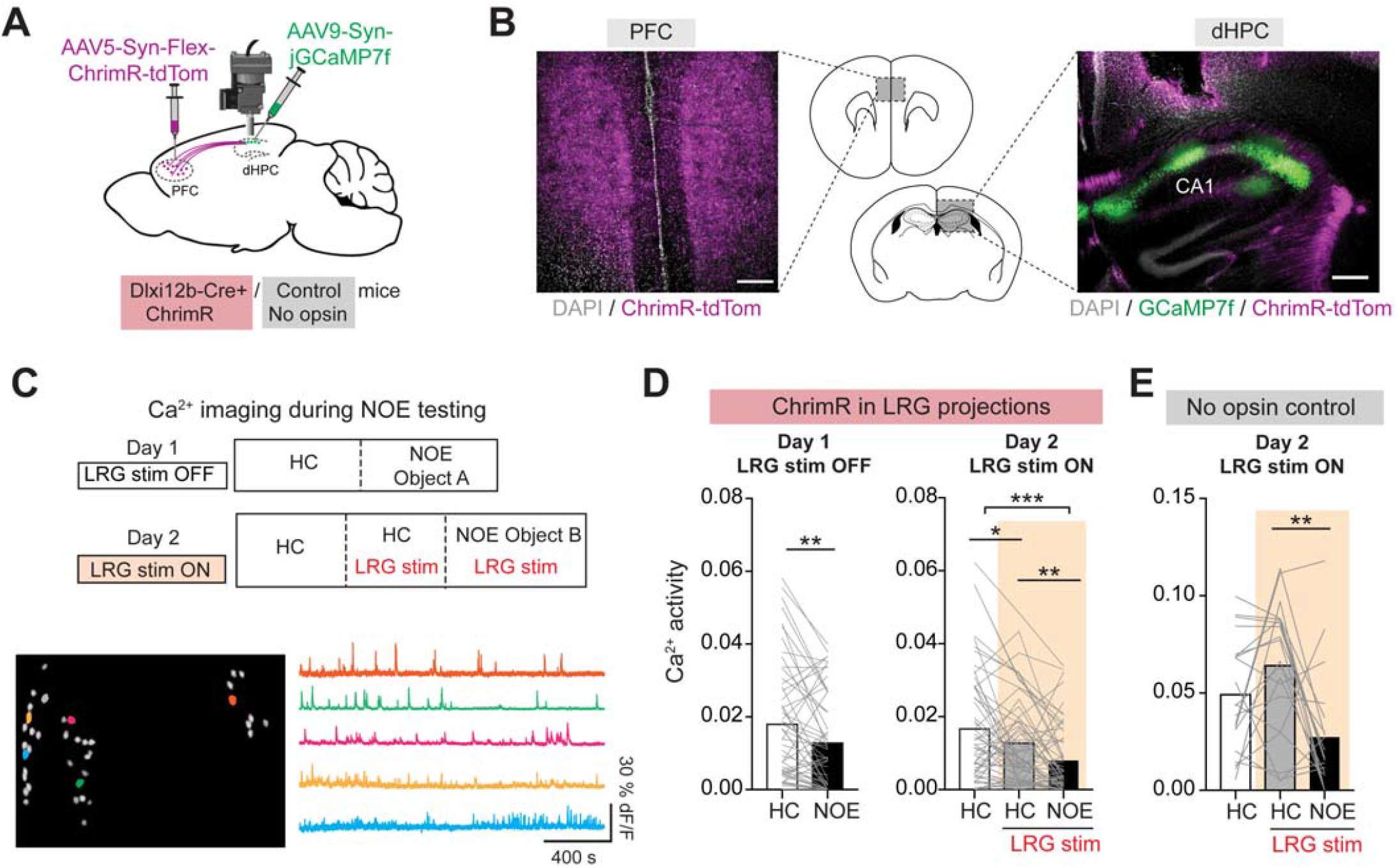
PFC–dHPC LRG projections shape CA1 neuronal activity during object exploration. **(A)** Strategy for *in vivo* Ca^2+^ imaging and optogenetic stimulation. Cre-dependent ChrimsonR-tdTomato (ChrimR-tdTom) virus was injected into the PFC of *Dlxi12b-Cre*+ and *Cre-negative* (control) mice; jGCamp7f virus was injected into dorsal CA1, and Ca^2+^ activity was imaged through an implanted GRIN lens connected to a miniscope. **(B)** Left: DAPI stained coronal section showing ChrimR-tdTom expression in PFC GABAergic neurons. Right: DAPI stained dHPC section showing jGCaMP7f expression in CA1 neurons and ChrimR-tdTom expression in PFC–dHPC LRG axonal fibers. Scale bars, 0.5 mm. **(C)** Top: on day 1, CA1 Ca^2+^ activity was measured during home cage (HC) and NOE epochs. On day 2, following an initial period of imaging in HC without optogenetic stimulation, Ca^2+^ imaging was combined with optogenetic stimulation of PFC–dHPC LRG projections (590–650 nm, ∼2 mW, 5 ms pulses at 20 Hz) during HC and NOE epochs. Bottom left: regions of interest (ROIs) corresponding to neurons from a representative Ca^2+^ imaging session. Bottom right: extracted dF/F Ca^2+^ transients from example dorsal CA1 neurons. Colors of the traces on the right correspond to the colored ROIs on the left. **(D)** Left: Ca^2+^ activity in CA1 neurons was significantly reduced during NOE. Each gray line represents a single neuron and bars represent the mean (n = 55 neurons from 2 mice); two-way paired t-test. Right: During the HC epoch on day 2, optogenetic stimulation of PFC–dHPC LRG projections reduced CA1 Ca^2+^ activity. On Day 2, activity was then further reduced during NOE (n = 59 neurons, 2 mice). One-way ANOVA followed by Tukey’s multiple comparison test; *** p < 0.001, ** p < 0.01, * p < 0.05. **(E)** Same as **D** for control (opsin-negative) mice. Light delivery alone did not affect Ca^2+^ activity. One-way ANOVA followed by Tukey’s multiple comparison test; ** p < 0.01. See also **Figure S6**.

### PFC–dHPC LRG projections enhance the encoding of objects by hippocampal ensembles

To assess whether these global changes in CA1 activity were associated with changes in how the hippocampus encodes NOE-relevant information, we compared the NOE-driven changes in neuronal activity on day 1 (no stimulation) vs. day 2 (LRG stimulation). As shown by the seminal discovery of place cells, dHPC CA1 neurons encode information by preferentially firing in specific spatial locations (Moser et al., 2008; O’Keefe, 1976; Wilson and McNaughton, 1993). Therefore, we asked whether the PFC–dHPC LRG projections affect the encoding of object location by individual hippocampal neurons. Specifically, for each neuron, we defined its ‘object signal-to-noise ratio’ (Object_SNR) as the change in its activity within a zone surrounding the object location before vs. after introducing the object (activity was z-scored relative to the mean and standard deviation outside the object zone) (Fig. 6A). Based on this metric, neurons that increased or decreased activity in the object zone by one standard deviation had Object_SNR of 1 or -1, respectively. During light stimulation, the activity of neurons decreased both within and outside of the object zone; the standard deviations of neuronal activity also decreased (Fig. 6B). Depending on exactly how these changes were distributed across neurons, Object_SNR values could potentially increase, decrease, or remain unchanged. In fact, we observed that stimulating PFC LRG projections significantly increased Object_SNR values relative to the no light condition (Fig. 6B, right-most panel). Notably, light delivery alone did not affect the neuronal activity or the Object_SNR in control (opsin-negative) mice (Fig. 6C). Furthermore, in opsin-expressing mice, LRG stimulation did not affect an analogous ‘SNR’ calculated for a control zone on the opposite side of the cage (instead of the object zone) (Fig. 6D, E). Thus, even though PFC–dHPC LRG projections potentiate feedforward inhibition and reduce overall network activity, their net effect on hippocampal encoding is to specifically enhance object-driven signals in individual CA1 neurons.

**Figure 6:**
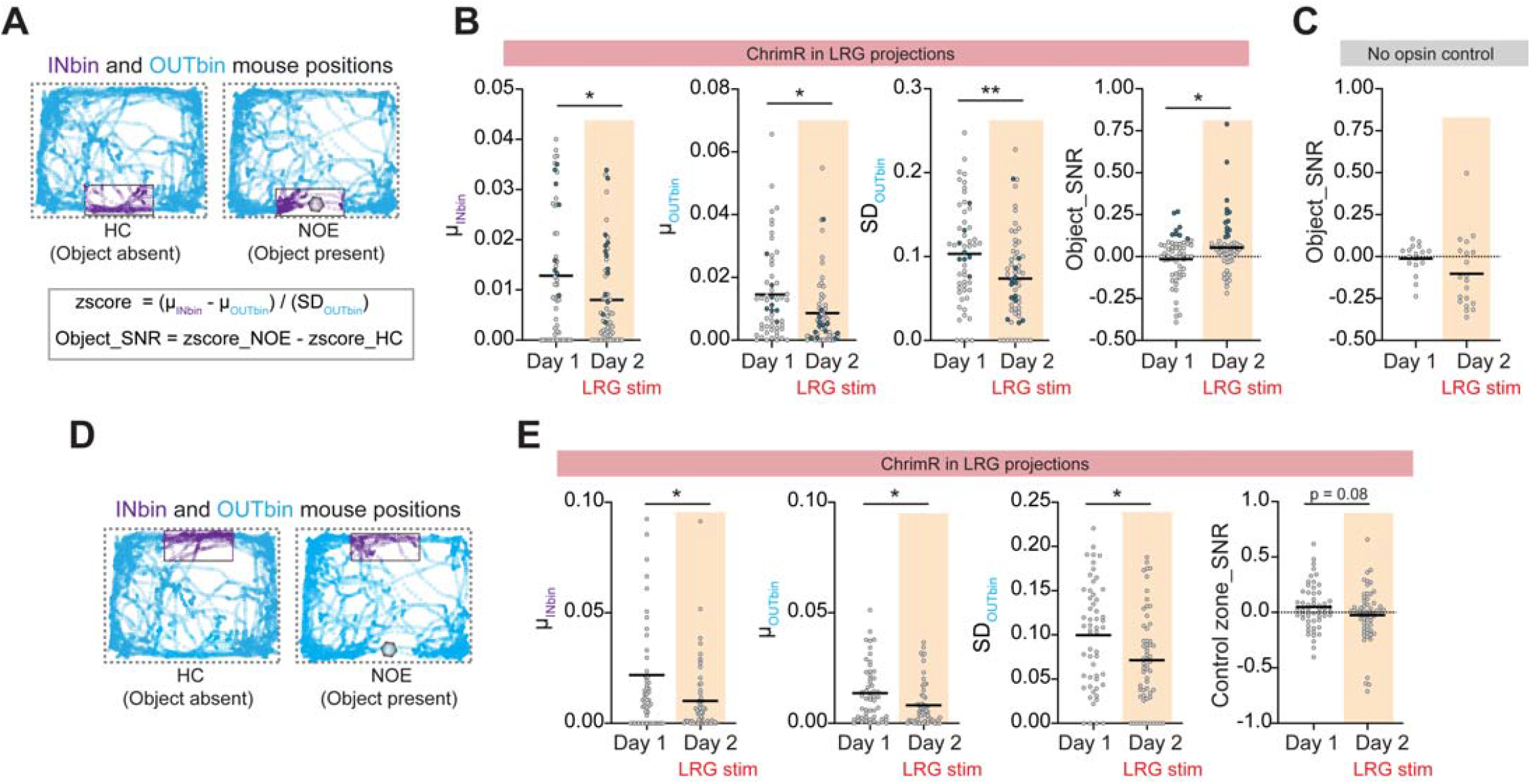
PFC–dHPC LRG projections increase neuronal signal-to-noise ratio for representation of object location in CA1. **(A)** Top: Frame-by-frame mouse positions during HC (left) and NOE (right) epochs are plotted. Example data from day 2 recording session is shown. Blue circles denote frames where mouse was outside the object zone (‘OUTbin’). Purple circles denote frames where mouse was in the object zone (‘INbin’). Gray shaded hexagon indicates the location of the novel object during NOE epoch and rectangle with black solid lines denotes the object zone. Bottom: z-scored Ca^2+^ activity was calculated as the difference between the mean Ca^2+^ activity when mouse was within (μ_INbin_) vs. outside (μ_OUTbin_) the object zone, divided by the standard deviation of activity outside the object zone (SD_OUTbin_). The difference in z-scored Ca^2+^ activity between HC to NOE epochs was used to compute Object_SNR. **(B)** Optogenetic stimulation of PFC–dHPC LRG projections significantly reduced μ_INbin_, μ_OUTbin_, and SD_OUTbin_. Object_SNR was increased on day 2 compared to day 1 in opsin-expressing mice. Empty gray circles represent from individual neurons and horizontal black lines show means. Filled blue circles indicate neurons exceeding an arbitrary threshold for Object_SNR (>0.1) to illustrate μ_INbin_, μ_OUTbin_, and SD_OUTbin_ for these high-SNR neurons. Two-way unpaired t-test; ** p < 0.01, * p < 0.05. **(C)** Object_SNR in control (opsin negative) mice was not affected by light delivery. **(D)** Frame-by-frame mouse positions during HC (left) and NOE (right) epochs are plotted. Example data from day 2 recording session is shown. Blue circles denote frames where mouse was away from a control zone (‘OUTbin’) that was the mirror image of the object zone, but on the opposite side of the cage. Purple circles denote frames where mouse was in the control zone (‘INbin’). Gray shaded hexagon denotes the novel object location in NOE epoch and rectangle with black solid lines denotes the control zone. **(E)** For the control zone, optogenetic stimulation of PFC–dHPC LRG projections significantly reduced μ_INbin_, μ_OUTbin_, and SD_OUTbin_ (similar to the object zone). However, SNR computed for the control zone was not affected by optogenetic stimulation of LRG projections (in contrast to the object zone). Empty gray circles represent values from individual neurons and black lines represent mean values. Two-way unpaired t-test; * p < 0.05.

## Discussion

Interactions between the PFC and HPC have been implicated in numerous aspects of cognition and emotion, including decisions about whether to engage in exploratory behaviors. While monosynaptic excitatory projections from the ventral HPC are believed to transmit specific information, e.g., the locations of goals, to the PFC (Spellman et al., 2015; Wang and Cai, 2006), pathways through which the PFC exerts top-down control over the HPC, and the exact nature of these top-down effects, have remained less well understood. Here, we describe a novel monosynaptic projection from the PFC-to-dHPC. There are many unusual features of this projection: it is GABAergic and targets hippocampal VIP+ ISIs, thus representing a ‘doubly disinhibitory’ long-range motif. We show that this projection modulates microcircuit dynamics in the CA1 region of dHPC, increasing feedforward inhibition, reducing spiking evoked by afferent inputs, and enhancing low-gamma activity that is synchronized between the PFC and dHPC. Furthermore, we show that activation of these projections reduces *in vivo* activity while specifically enhancing the representations of objects in the dorsal CA1. Lastly, in accord with the postulated role of PFC as a top-down controller, we found that the activation of these LRG projections drives object exploration behavior in mice. Overall, our study shows that these top-down prefrontal projections can dynamically control the network state and emergent circuit function in the dHPC, thereby altering the signal-to-noise ratio for specific neural representations and eliciting corresponding changes in behavior. This answers long-standing questions about the mechanisms and nature of top-down control in the limbic system.

### Relationship to previous work

Multiple lines of evidence in humans, non-human primates, and rodents have suggested that PFC can exert top-down control over information processing in the HPC, particularly during behaviors involving spatial and object exploration (Brincat and Miller, 2015; Eichenbaum, 2017; Jin and Maren, 2015; Place et al., 2016; Preston and Eichenbaum, 2013; Shin and Jadhav, 2016; Sigurdsson and Duvarci, 2016; Yu and Frank, 2015). Importantly, previous studies have shown that lesion and pharmacological inactivation of PFC severely impairs spatial navigation and object exploratory behaviors, and also disrupts neuronal encoding in the HPC (Churchwell et al., 2010; DeVito and Eichenbaum, 2010; Floresco et al., 1997; Guise and Shapiro, 2017; Kyd and Bilkey, 2003; Wang and Cai, 2006; Yoon et al., 2008). Nevertheless basic aspects of this process have remained elusive. Specifically, the pathways mediating prefrontal top-down control have not been identified, and it was not known whether the PFC acts by transmitting specific information to the HPC vs. by modulating the network state and emergent circuit function. Addressing these questions is crucially important, because interactions between the HPC and PFC have been implicated in so many behaviors and disorders, and because the role of the PFC in top-down control is largely taken for granted despite the paucity of knowledge about specific mechanisms. Our study addresses this major gap by revealing a novel anatomical substrate mediating prefrontal top-down control, and showing exactly how it regulates behavior via actions on hippocampal neurons, microcircuits, network dynamics, and information processing.

Our anterograde tracing experiments showed that the PFC LRG projections are concentrated in the dorsal HPC, relative to the intermediate and ventral parts portions of HPC. By contrast, the NR, which is known to mediate indirect PFC–HPC communication, preferentially innervates the intermediate and ventral HPC (Hoover and Vertes, 2012). This suggests that conjunctive information transfer via the direct PFC LRG pathway and the indirect PFC→NR→HPC pathway would allow PFC to orchestrate activity along the entire extent of the hippocampal dorsoventral axis. Interestingly, the hippocampal dorsoventral axis is functionally segregated with the dorsal HPC being crucially involved in spatial processing and the ventral HPC regulating emotions, fear, and anxiety (Fanselow and Dong, 2010). Therefore, an alternate possibility is that the direct PFC→dHPC LRG projections and indirect PFC→NR→ventral HPC projections may mediate fundamentally distinct aspects of top-down control over cognitive vs. emotional behaviors, respectively. Similar to the functional segregation along the hippocampal dorsoventral axis, the PFC can also be subdivided dorsoventrally, into functionally specialized subregions, e.g., anterior cingulate (ACC), prelimbic (PL), and infralimbic (IL) cortices. We found PFC–dHPC LRG projections originating from PL. Notably, the ACC also sends direct projections to the dHPC (Rajasethupathy et al., 2015). However, by targeting excitatory neurons in the CA3 subregion of the HPC, these previously described ACC–CA3 excitatory projections primarily regulate the retrieval of fear memories. While the role of ACC–CA3 projections in spatial exploratory behaviors has not been investigated, it is plausible that the direct projections originating from ACC and PL regions transmit parallel streams of information from the PFC to alter hippocampal activity during distinct behaviors. Future work will be necessary to elucidate how glutamatergic vs. GABAergic top-down projections originating from different regions of the PFC, and targeting different portions of the HPC potentially interact and/or complement each other.

Our study shows that PFC–dHPC LRG projections target inhibitory interneurons, but not excitatory pyramidal neurons, within the CA1 subregion. This preferential targeting of inhibitory interneurons is similar to what has been observed in previous studies of cortical LRG inputs to the HPC. Specifically, the entorhinal cortex, which is the primary interface between the HPC and the neocortex, sends LRG projections which target local interneurons in the HPC (Basu et al., 2016; Melzer et al., 2012). However, in contrast to effect we observed, whereby PFC–dHPC LRG projections increase feedforward inhibition by inhibiting VIP interneurons, entorhinal LRG projections primarily act to reduce hippocampal feedforward inhibition. This raises the possibility that feedforward inhibition may be a convergent pathway on which many LRG inputs act to regulate hippocampal information processing. Feedforward inhibition represents an attractive target, as it crucially regulates input-output gain, neuronal plasticity, and information encoding in hippocampal pyramidal neurons (Grienberger et al., 2017; McKenzie, 2018).

VIP interneurons in the HPC are a specialized class which disinhibit other GABAergic interneurons, thereby tending to promote increases in microcircuit activity. Accordingly, disinhibition mediated by hippocampal VIP interneurons has been implicated in gain control, memory, selective attention, and goal-directed behaviors (Cunha-Reis and Caulino-Rocha, 2020; Turi et al., 2019). While we found that the CA1 interneurons which receive PFC–dHPC LRG inputs are electrophysiologically heterogeneous but tend to express VIP, understanding whether their axons target specific inhibitory loci within the CA1 microcircuit will help us to further understand the detailed nature of their actions. This is important because VIP+ interneurons in the CA1 subregion constitute electrophysiologically and morphologically diverse subtypes (Acsády et al., 1996a; 1996b; Chamberland and Topolnik, 2012). Prior work has also shown that specialized subpopulations of hippocampal VIP+ GABAergic neurons send long-range projections which innervate different parts of the hippocampal formation, and are recruited during specific oscillatory states (Francavilla et al., 2018). It is possible that PFC–dHPC LRG projections target these long-range projecting VIP neurons, contributing to their state-dependent patterns of activity, and helping to produce some of the changes in network dynamics we observed here. Alternatively, the changes in hippocampal low-gamma oscillations we observed could be result from increases in feedforward inhibition.

### Relationship to disease

Many neuropsychiatric disorders – including autism, schizophrenia, depression, and anxiety disorders – are proposed to involve deficits in prefrontal functions and top-down control (Gilbert et al., 2008; Hare and Duman, 2020; Orellana and Slachevsky, 2013), and in particular, altered connectivity and communication between PFC and HPC (Cunniff et al., 2020; Godsil et al., 2013; Kupferschmidt and Gordon, 2018; Li et al., 2015; Sigurdsson et al., 2010). Abnormalities in GABAergic neuron structure and function have also been heavily implicated in the pathophysiology of neuropsychiatric disorders (Chattopadhyaya and Cristo, 2012; Marin, 2012; Paterno et al., 2020). PFC–dHPC LRG projections obviously represent a point of convergence for these different mechanisms. Thus, abnormalities in PFC–dHPC LRG projections could plausibly contribute to the disruptions in network oscillations, top-down control, and PFC-dHPC communication that occur in a variety of disease states.

## Conclusion

In summary, our study describes a novel anatomical pathway which plays a key role in direct PFC-to-HPC communication. The unique features of these projections (i.e., long-range GABAergic, disinhibitory interneuron targeting) enable PFC to dynamically alter emergent network activity and information processing in the HPC, and thereby exert top-down control over exploratory behavior.

## Acknowledgements

We thank M. Sharma and H. Seifikar for technical support. We are grateful to L. Frank, M. Kheirbek, and members of the Sohal laboratory for comments on earlier versions of this manuscript. This work was supported by National Institute of Mental Health (R01MH106507 and RO1MH117961 to V.S.S.) and 2018 NARSAD Young Investigator Grant from Brain & Behavior Research Foundation (Leichtung Family investigator, R.M.). Confocal images were acquired at the Nikon Imaging Center at the University of California San Francisco.

## Author contributions

R.M. and V.S.S. designed the experiments and analyses. R.M. performed all experiments and analyzed the data, except that R.M. and Y.L. performed immunohistochemistry. S.S. generated pilot histology data for anterograde tracing experiments. R.M. and V.S.S. wrote the manuscript.

## METHODS

### Animals

All animal care procedures and experiments were conducted in accordance with the National Institutes of Health guidelines and approved by the Administrative Panels on Laboratory Animal Care at the University of California, San Francisco. Mice were housed in a temperature-controlled environment (22–24 °C) with ad libitum access to food and water. Mice were reared in normal lighting conditions (12-h light/dark cycle). Adult mice from the following lines were used: *Dlxi12b-Cre* (Potter et al., 2009) and wild-type CD-1.

### Virus and retrograde tracer injections

Mice were anesthetized with isoflurane and placed on a stereotaxic frame (David Kopf Instruments). An incision was made to expose the skull, and bregma and lambda were used as references to align the skull. Body temperature was maintained using a heating pad. Virus was injected (at the rate of 100 nl/min) with a microinjection syringe (Nanofil 10 µl with 35 gauge needle, World Precision Instruments) connected to a microsyringe pump (World Precision Instruments, UMP3 UltraMicroPump). Coordinates for injections into PFC were (in mm, relative to Bregma) 1.8 anterior-posterior (AP), ±0.3 mediolateral (ML), -2.4 dorsoventral (DV); and coordinates for injections into dHPC were -1.35 AP, ±0.65 ML, -1.5 DV.

For anterograde tracing, ChR2 assisted circuit-mapping and optogenetic stimulation experiments, either AAV5-EF1α-DIO-eYFP virus or AAV5-EF1α-DIO-ChR2-eYFP virus (UNC Vector core, 650 nl) was injected into PFC of *Dlxi12b-Cre*+ mice and *Cre-negative* mice. For intersectional labeling of dHPC projecting PFC LRG neurons, CAV2-Cre virus (del Rio et al., 2019; Hnasko et al., 2006) (650 nl) was injected in the dHPC and AAV5-Dlxi12b-BG-DIO-ChR2-eYFP virus (Lee et al., 2014) (650 nl) was injected in the PFC of CD-1 mice. For retrograde labeling of dHPC projecting PFC LRG neurons, Alexa Flour 594 Cholera toxin beta subunit conjugate (CTb-594, Invitrogen; 0.5% w/v, 400–500 nl) was injected in dHPC of CD-1 mice. After virus or tracer injection, the microinjector needle was left in place for 5–6 min before being removed from the brain. Mice were sutured (if receiving viral/tracer injection only) and were allowed to recover on a heated pad until ambulatory.

### Optic fiber implantation

Following bilateral AAV5-EF1α-DIO-ChR2-eYFP virus injection in PFC, dual fiber-optic cannulas (Doric lenses; 200/240 mm, 0.22NA) were implanted in dHPC (−1.35 AP, ±0.65 ML, -1.4 DV). During these surgeries, the skull was scored with a scalpel to improve implant adhesion. We waited at least 7 weeks after surgery to allow time for viral expression.

### Optogenetic stimulation of PFC–dHPC LRG projections

A 473 nm blue laser (OEM Laser Systems, Inc.) was coupled to the dual fiber-optic cannula (implanted in dHPC) through a dual fiber-optic patch cord (Doric Lenses, Inc.), and was controlled via a function generator (Agilent 33500B Series Waveform Generator). Laser power was adjusted such that the final light power was 3–4 mW total, summed across both fibers, and averaged over 20 Hz light pulses (5 ms duration).

### Behavioral assays

After sufficient time for surgical recovery and viral expression, mice were handled and habituated for multiple days (3–5 days). Briefly, mice were first habituated to the behavioral testing room for 30 min prior to handling each day. For 2–3 days before starting testing, mice were habituated to the cable tethers in their home cage for 15 min. The experimenter was blinded to experimental groups during behavioral testing and scoring. A USB webcam (Logitech) connected to a computer running ANY-maze (Stoelting Co.) was used to record behavior movies. The position of mice was tracked using the built-in tracking in ANY-maze software. In some experiments, mouse positions were tracked using trained neural networks in DeepLabCut open-source software package (Mathis et al., 2018).

#### Novel object exploration

For measuring novel object exploration, one previously unexplored object was placed in the home cage of an experimental mouse for 5 min. A blinded observer manually scored the following parameters: exploration time, bouts of exploration, and latency to the first exploration. Objects used in our study were usually lego toys, dice, small plumbing connectors, and falcon tube caps. For experimental mice with dual-fiber optic implants, two object interaction tests were performed over two days: day 1 testing was performed without light stimulation, and day 2 testing was done during optogenetic stimulation of PFC–dHPC LRG projections. *Cre-negative* mice (no opsin control) with dual-fiber optic implants underwent similar behavior testing procedures.

#### Social interaction test

For social interaction test, a novel juvenile (3–4 week old) mouse of the same sex was introduced in the home cage of an experimental mouse for 5 min. A blinded observer manually scored the time (in seconds) the experimental mouse spent with its nose in direct contact with the novel juvenile intruder. For all experimental mice, two social interaction tests were performed over two days: day 1 testing was performed without light stimulation, and day 2 testing was done during optogenetic stimulation of PFC LRG projections.

#### Open field exploration test

Mice were placed in the center of a 50 x 50 cm open-field arena and were allowed to freely explore for 12 min. The testing time was divided into four (3 min) epochs. PFC–dHPC LRG projections were optogenetically stimulated during the 2^nd^ and 4^th^ epochs. Distance traveled during no stimulation (light OFF) and during optogenetic stimulation (light ON) epochs was quantified using the ANY-maze tracking software.

#### Real-time place preference (RTPP) test

Real-time place preference (RTPP) testing protocol consisted of three 20 min sessions conducted over 3 days. An apparatus with two identical chambers was used for RTPP testing. On day 1, mice were habituated to the apparatus for 15 min. On day 2, mice were placed into one randomly chosen chamber and the time spent in the two chambers was recorded. On day 3, one of the chambers was randomly assigned as the stimulated chamber. When mice entered this chamber, they received 20 Hz laser pulses (473 nm, 3–4 mW, 5 ms). The ratio of the time spent in the simulated chamber vs. the non-stimulated chamber was used as the preference index. The sides of the stimulated chambers were counterbalanced across all mice.

### LFP recordings: surgery and analysis

#### Surgery

Mice were anesthetized with isoflurane and placed on a stereotactic frame. After cleaning, the skull was scored with a scalpel to improve implant adhesion. For LFP recordings from wild-type CD-1 mice, tungsten electrodes (Microprobes) were inserted into the PFC (1.8 AP, -0.3 ML, -2.4 DV) and dHPC (−1.35 AP, -0.65 ML, -1.5 DV). For multisite LFP recordings combined with optogenetics, one LFP electrode was implanted after AAV5-EF1α-DIO-ChR2-eYFP virus injection into PFC of *Dlxi12b-Cre*+ mice. A custom-made optrode (optical fiber + electrode) was implanted in dHPC to stimulate ChR2+ PFC–dHPC LRG axon terminals during LFP recordings. To fabricate optrodes, a tungsten LFP recording electrode was affixed to one of the fibers of the dual-fiber optic cannula such that the tip of the electrode protruded 200–300 µm beyond the end the optic fiber (Lee et al., 2019). Reference and ground screws were implanted above the cerebellum. Electrodes and screws were cemented to the skull with Metabond (Parkell) and connected to a headstage for multi-channel recordings (Pinnacle). Following surgery, mice were monitored postoperatively, given analgesics, and individually housed.

#### Recording and analysis

LFP data were acquired at 2 KHz and band-pass filtered from 0.5-150Hz. Electrode placements were histologically confirmed. Analysis of LFP data was done using custom MATLAB (Mathworks) scripts. Briefly, signals were imported into MATLAB and LFP log power (for both channels) was calculated using the power spectral density output from the spectrogram function. For phase-synchrony and amplitude covariance analysis, LFPs were FIR-filtered for different frequency bands, then Hilbert transformed to yield the instantaneous amplitudes and phases. The following frequency bands were compared: theta band (4–12 Hz), beta band (15–25 Hz), low-gamma band (25–55 Hz), and high-gamma band (65–85 Hz).

To detect nonzero phase interdependencies (phase synchrony) between LFP signals recorded at PFC and dHPC electrodes, we estimated the weighted Phase Lag Index (wPLI) (Vinck et al., 2011) using the imaginary component of the cross-spectrum (S_xy_) (Equations 1.1 and 1.2). A_x_ and A_y_ are instantaneous amplitudes; and Φx and Φy are instantaneous phases for PFC and dHPC signals, respectively.

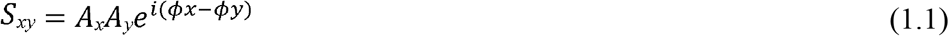

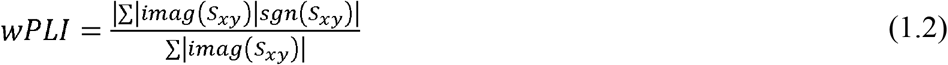

The amplitude covariation between PFC and dHPC was calculated as the maximum normalized cross-correlation (xcorr function in MATLAB) of the instantaneous band-pass filtered amplitudes of LFP signals at each electrode. Coherence between LFP signals was computed using the mscohere function in MATLAB. Log power, wPLI, amplitude covariation, and coherence were calculated over short time intervals (at least 3 sec in duration), i.e., the intervals during which a mouse was actively exploring an object or matched intervals during baseline periods when the mouse was in its home cage.

### *Ex-vivo* slice physiology

#### Slice preparation

Adult mice were anesthetized with an intraperitoneal injection of euthasol and transcardially perfused with an ice-cold cutting solution containing (in mM) 210 sucrose, 2.5 KCl, 1.25 NaH_2_PO_4_, 25 NaHCO_3_, 0.5 CaCl_2_, 7 MgCl_2_, 7 dextrose (bubbled with 95% O_2_–5% CO_2_, Ph ∼7.4). Mice were decapitated and the brains were removed. For acute prefrontal sections: two parallel cuts were made along the coronal plane at the rostral and caudal ends of the brains; brains were mounted on the flat surface created at the caudal end; three coronal slices (250□µm thick) were obtained using a vibrating blade microtome (VT1200S, Leica Microsystems Inc.). Dorsal hippocampal (dHPC) slices were obtained using a blocking technique described previously (Malik et al., 2015). Briefly, dHPC slices were obtained by making a blocking cut at a 45° angle from the coronal plane starting at the posterior end of the forebrain. A second blocking cut was made at 45° relative to the coronal plane, but starting from approximately one-third of the total length of the forebrain (from the most anterior point). Brains were mounted on the flat surface created by the first blocking cut. Approximately, 3 dorsal slices were obtained from each hemisphere.

Slices were allowed to recover at 34□°C for 30□min followed by 30□min recovery at room temperature in a holding solution containing (in mM) 125 NaCl, 2.5 KCl, 1.25 NaH_2_PO_4_, 25 NaHCO_3_, 2 CaCl_2_, 2 MgCl_2_, 12.5 dextrose, 1.3 ascorbic acid, 3 sodium pyruvate.

#### Ex-vivo patch clamp recordings

Somatic whole-cell current-clamp and voltage-clamp recordings were obtained as previously described (Malik and Johnston, 2017; Malik et al., 2019). Briefly, submerged slices were perfused in heated (32–34□°C) artificial cerebrospinal fluid (aCSF) containing (in mM): 125 NaCl, 3 KCl, 1.25 NaH_2_ PO_4_, 25 NaHCO_3_, 2 CaCl_2_, 1 MgCl_2_, 12.5 dextrose (bubbled with 95% O_2_/5% CO_2_, pH ∼7.4). Neurons were visualized using DIC optics (and eYFP fluorescence in a few experiments) fitted with a 40x water-immersion objective (BX51WI, Olympus microscope). During recordings from prefrontal slices, dHPC projecting PFC LRG neurons in all cortical layers were identified by eYFP expression. During recordings from hippocampal slices, CA1 pyramidal neurons (PNs) and CA1 local inhibitory neurons (INs) were identified using laminar location (under DIC optics) and intrinsic properties of the recorded neurons.

Patch electrodes (2–3□MΩ) were pulled from borosilicate capillary glass of external diameter 1□mm (Sutter Instruments) using a Flaming/Brown micropipette puller (model P-2000, Sutter Instruments). For current-clamp recordings, electrodes were filled with an internal solution containing the following (in mM): 134 K-gluconate, 6 KCl, 10 HEPES, 4 NaCl, 7 K_2_-phosphocreatine, 0.3 Na-GTP, and 4 Mg-ATP (pH ∼7.3 adjusted with KOH). Biocytin (Vector Laboratories) was included (0.1–0.2%) for subsequent histological processing of recipient CA1 neurons. For voltage-clamp recordings, the internal solution contained the following (in mM): 130 Cs-methanesulfonate, 10 CsCl, 10 HEPES, 4 NaCl, 7 phosphocreatine, 0.3 Na-GTP, 4 Mg-ATP, and 2 QX314-Br (pH ∼;7.3 adjusted with CsOH). In a few recordings, 15 μm AlexaFluor-594 (Invitrogen) was also added to the internal solution. Electrophysiology data were recorded using Multiclamp 700B amplifier (Molecular Devices). Voltages have not been corrected for measured liquid junction potential (∼8□mV). Data collection was started 5–8 min after successful transition to the whole-cell configuration. Series resistance and pipette capacitance were appropriately compensated before each recording. Series resistance was usually 10– 20□MΩ, and experiments were terminated if series resistances exceeded 25□MΩ.

#### Data analysis

*Ex-vivo* electrophysiology data were analyzed using custom routines written in IGOR Pro (Wavemetrics). Resting membrane potential (RMP) was measured in current-clamp mode immediately after reaching whole-cell configuration. Input resistance (Rin) was calculated as the slope of the linear fit of the voltage-current plot generated from a family of hyperpolarizing and depolarizing current injections (−50 to +20 pA, steps of 10□pA). Firing output was calculated as the number of action potentials (APs) fired in response to 800□ms long depolarizing current injections (25–500□pA). Firing frequency was calculated as the number of APs fired per second. Firing traces in response to 50 pA current above the rheobase were used for analysis of single AP properties – AP threshold, maximum *dV*/*dt* (rate of rise of AP), AP amplitude, AP half-width, and fast afterhyperpolarization (fAHP) amplitude. AP threshold was defined as the voltage at which the value of third derivative of voltage with time is maximum. Action potential amplitude was measured from threshold to peak, and the half-width was measured at half this distance. Fast afterhyperpolarization was measured from the threshold to the negative voltage peak after the AP. Index of spike-frequency accommodation (SFA) was calculated as the ratio of the last inter-spike interval to the first inter-spike interval.

Recorded inhibitory neurons (INs) in PFC and dorsal CA1 were classified as fast spiking, regular spiking or irregular spiking based on electrophysiological properties. Specifically, INs were classified as fast spiking if they met 3 out of the 4 following criteria: AP half-width was < 0.5□ms, firing frequency > 50□Hz, fAHP amplitude >14□mV, and SFA index < 2. Irregular spiking INs were initially visually identified based on their high variability in inter-spike interval and burst-like intermittent spiking properties. This classification was confirmed using a firing frequency threshold (<50 Hz) and/or a SFA index threshold (>2). Dorsal CA1 neurons were classified as pyramidal neurons if they satisfied the following criteria: cell body located in stratum pyramidale, AP half-width > 1 ms, fAHP amplitude < 5 mV, and maximum firing frequency < 20 Hz.

To measure optogenetically evoked spiking in ChR2-eYFP+ PFC INs and to measure optogenetically evoked postsynaptic currents (oPSCs) in CA1 neurons, ChR2 was stimulated using 5□ms long light pulses (maximum light power, 4□mW/mm^2^) generated by a Lambda DG-4 high-speed optical switch with a 300□W Xenon lamp (Sutter Instruments) and an excitation filter centered around 470□nm. Light pulses were delivered to the slice through a 40x objective (Olympus). To measure the reversal potential of oPSCs, the holding potentials were systematically varied from −100 to +20 mV in 10 mV steps. The drugs applied were 6-cyano-7-nitroquinoxaline-2,3-dione disodium salt hydrate (CNQX), 2-(3-carboxypropyl)-3-amino-6-(4 methoxyphenyl)-pyridazinium bromide (Gabazine), and d-2-amino-5-phosphonopentanoic acid (D-AP5) (Tocris). Drugs were prepared as concentrated stock solutions and were diluted in ACSF on the day of the experiment.

To measure afferent input mediated feedforward excitation and inhibition in CA1 PNs, bipolar stimulating electrodes (Microprobes) were placed at stratum radiatum (SR) and stratum lacunosum-moleculare (SLM) to stimulate Schaffer collateral (SC) and temporo ammonic (TA) inputs, respectively. The protocol for theta-burst stimulation (TBS) consisted of bursts with five electrical stimulations (40 Hz) repeated at 5 Hz. To measure the effect of PFC LRG inputs on firing output and EPSP summation during TBS protocol, train of 470 nm light pulses (20 Hz, 5 ms) was delivered through the 40x objective. Firing frequency during TBS was calculated as the average number or APs fired per burst, and summation was estimated as the area of the last EPSP in the TBS train.

### *In vivo* Ca^2+^ imaging

#### Surgery

Mice underwent two stereotactic surgeries. Cre-dependent AAV5-Syn-FLEX-ChrimsonR-tdTomato virus (Addgene) was injected in PFC (1.8 AP, ±0.3 ML, -2.4 DV) of *Dlxi12b-Cre*+ and *Cre-negative* mice. Following this, 500–550 nl of AAV9-Syn-jGCaMP7f-WPRE virus (diluted 1:2; Addgene) was injected in dorsal CA1 to express synapsin-driven calcium sensor jGCaMP7f (injection coordinates: -1.4 AP, +0.8 ML, -1.5 DV). After 3–4 weeks of viral expression, cortex overlying dorsal CA1 was slowly aspirated and a 1 mm diameter x 4 mm long integrated GRIN lens (Inscopix) was slowly advanced above the dorsal CA1 and cemented in place with Metabond dental cement. Mice were allowed to recover for at least 3 weeks before starting behavior and imaging experiments.

#### Combined Ca^2+^ imaging and optogenetics

Imaging data were collected using a miniaturized one-photon microscope (nVoke2; Inscopix Inc.). GCaMP7f signals (Ca^2+^ activity) were detected using 435–460 nm excitation LED (0.1–0.2 mW), and optogenetic stimulation of ChrimR expressing axons was performed using a second excitation LED centered around 590–650 nm (5 ms pulses at 20 Hz, 1–2 mW light power). Ca^2+^ movies were acquired at 20 frames per second, spatially downsampled (4x), and were stored for offline data processing.

Mice were placed into a large housing cage (48 x 35 cm) for 2–3 days for 20 min where they habituated to the scope. After habituation, mice underwent a two-day behavioral testing protocol for recording NOE related Ca^2+^ activity in CA1 neurons. On day 1, mice were allowed to explore the large home cage for 15 min (HC epoch). Following this, mice were allowed to explore a novel object introduced in the cage for 15 min (NOE epoch). On day 2, mice were allowed to explore the home cage for 15 min (HC epoch) followed by optogenetic stimulation during home cage exploration for 10 min (HC + LRG stim epoch). Mice were then allowed to explore a novel object combined with optogenetic stimulation of PFC–dHPC LRG projections (NOE + LRG stim epoch). The behavior of mice during different epochs was recorded using ANY-maze software, and input TTL pulses from ANY-maze to nVoke2 acquisition software were used to synchronize Ca^2+^ imaging and mouse behavior movies.

#### Data analysis

Ca^2+^ imaging movies were preprocessed using Inscopix Data Processing Software (IDPS; Inscopix, Inc.). The video frames were spatially filtered (band-pass) with cut-offs set to 0.005 pixel^-1^ (low) and 0.5 pixel^-1^ (high) followed by frame-by-frame motion correction for removing movement artifacts associated with respiration and head-unrestrained behavior. The mean image over the imaging session was computed, and the dF/F was computed using this mean image. The resultant preprocessed movies were then exported into MATLAB, and cell segmentation was performed using an open-source calcium imaging software (CIAPKG) (Corder et al., 2019). Specifically, we used a Principal Component Analysis/Independent Component Analysis (PCA/ICA) approach to detect and extract ROIs (presumed neurons) per field of view (Mukamel et al., 2009). For each movie, the extracted output neurons were then manually sorted to remove overlapping neurons, neurons with low SNR, and neurons with aberrant shapes.

Accepted neurons and their Ca^2+^ activity traces were exported to MATLAB for further analysis using custom scripts (as previously described in (Frost et al., 2020)). Briefly, we calculated the standard deviation (σ) of the Ca^2+^ movie and used this to perform threshold-based event detection on the traces by first detecting increases in dF/F exceeding 2σ (over one second). Subsequently, we detected events that exceeded 10σ for over two seconds and had a total area under the curve higher than 150σ. The peak of the event was estimated as the local maximum of the entire event. For an extracted output neuron, active frames were marked as the period from the beginning of an event until the Ca^2+^ signal decreased 30% from the peak of the event (up to a maximum of 2 seconds).

#### Procedure for measuring object-related changes in Ca^2+^ activity

Frame-by-frame x-y positions of the head of a mouse in the testing cage were detected using DeepLabCut. A small rectangular area surrounding the object location in the testing cage was marked as the object zone. Time points (frames acquired at 30 Hz and resampled at 20 Hz, using resample function in MATLAB) when the mouse’s head was inside the object zone were classified as INbin and the remaining frames were classified as OUTbin. We then recorded the frame-by-frame *Ca*^*2*+^ *activity* of neurons corresponding to the INbin and OUTbin position frames. For all extracted neurons, the mean activity for INbin (μ_INbin_) and OUTbin (μ_OUTbin_) frames were calculated. We also calculated the standard deviation (SD_OUTbin_) of neuronal activity in OUTbin frames. The z-scored activity of each neuron was estimated using equation 2.1. The object signal-to-noise ratio (Object SNR) was calculated using the z-scored activity during HC and NOE epochs (Equation 2.2).

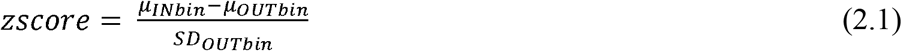

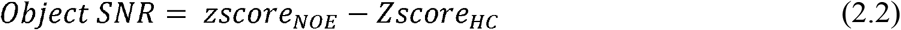

### Histological processing

#### Assessment of virus expression and anterograde tracing of LRG projections

Animals were transcardially perfused with PBS, and then with 4% paraformaldehyde (PFA). The brains were post-fixed for at least one day in PFA solution. Coronal sections (50–75 µm thick) were obtained using a vibratome. Sections that included the injection sites, electrode implantation sites and lens implantation sites were mounted on slides and cover-slipped using a glycerol-base, aqueous mounting medium (Vectashield Plus Antifade Mounting Medium, Vector labs). Sections were first scanned using an upright wide-field fluorescence microscope. Following this, confocal images were taken with 10x and 20x objectives on an Andor Borealis CSU-W1 spinning disk confocal mounted on a Nikon Ti Microscope (UCSF Nikon Imaging Center, NIH S10 Shared Instrumentation grant 1S10OD017993-01A1) and captured with an Andor Zyla sCMOS camera and Micro-Manager software (Open Imaging).

#### Inhibitory neuron marker expression in recipient CA1 neurons

Slices containing biocytin-filled cells were fixed overnight in a buffered solution containing 4% PFA. Slices were rinsed in PBS, then blocked and permeabilized in PBS with 5% Donkey Serum, 0.3% Triton X-100 and 1% BSA. Slices were immuno-stained overnight with one or two primary antibodies: rabbit anti-PV (Swant; diluted 1:200), rat anti-SST (Millipore, diluted 1:200), or rabbit anti-VIP (Immunostar, diluted 1:200). Slices were washed 6 x 10min in PBS containing 0.3% Triton X-100. Slices were incubated with donkey anti-rabbit Alexa-488, donkey anti-rat Alexa 594 secondary antibody (1:800, Thermo Fisher), and Streptavidin-647 (1:300, Thermo Fisher) overnight at 4°C. After washing 6 x 10min in PBS with 0.3% Triton X-100, slices were mounted with an aqueous mounting medium. Confocal mages were obtained as described above.

#### Inhibitory neuron (IN) marker expression in CTb tagged PFC LRG neurons

5-7 days after CTb injection, mice were transcardially perfused with PBS followed by 4% PFA solution, and brains were post-fixed for at least one day. Coronal sections (75 µm) were obtained using a vibratome, and immuhistochemistry was performed (as described above). The following primary antibodies were used to stain for IN markers: rabbit anti-PV (Swant; diluted 1:200); rat anti-SST (Millipore, diluted 1:200); rabbit anti-VIP (Immunostar, diluted 1:200); rabbit anti-NPY (Immunostar, diluted 1:500); rabbit anti-calretinin (Immunostar, diluted 1:500); rabbit anti-nNOS (Life technologies, diluted 1:500), and goat anti-CTb (List, diluted 1:500). The following secondary antibodies were used: donkey anti-rabbit Alexa 488; donkey anti-rat Alexa 488; and donkey anti-goat Alexa 594. For each IN marker, confocal images collected from mounted sections were used to manually count the number of CTb+ and IN marker+ PFC neurons (ImageJ software).

### Statistical analysis

Detailed statistical analyses were performed using MATLAB and Graphpad Prism. Comparisons of means were performed using paired or unpaired two-tailed Student’s t test, one-way ANOVA or two-way repeated measures ANOVA with Tukey post hoc test unless otherwise stated. For non-parametric data sets, we used a Chi-square test to determine significance. Sample sizes and statistical tests and parameters are listed in the figure legends. Data are reported as mean ± S.E.M. unless otherwise stated.

## Data availability

Data supporting the findings of this study are available from the corresponding author upon reasonable request.

## Code availability

Custom code used in this study is available from the corresponding author upon reasonable request.

## SUPPLEMENTAL INFORMATION: FIGURES AND LEGENDS

**Figure S1, related to Figure 1:**
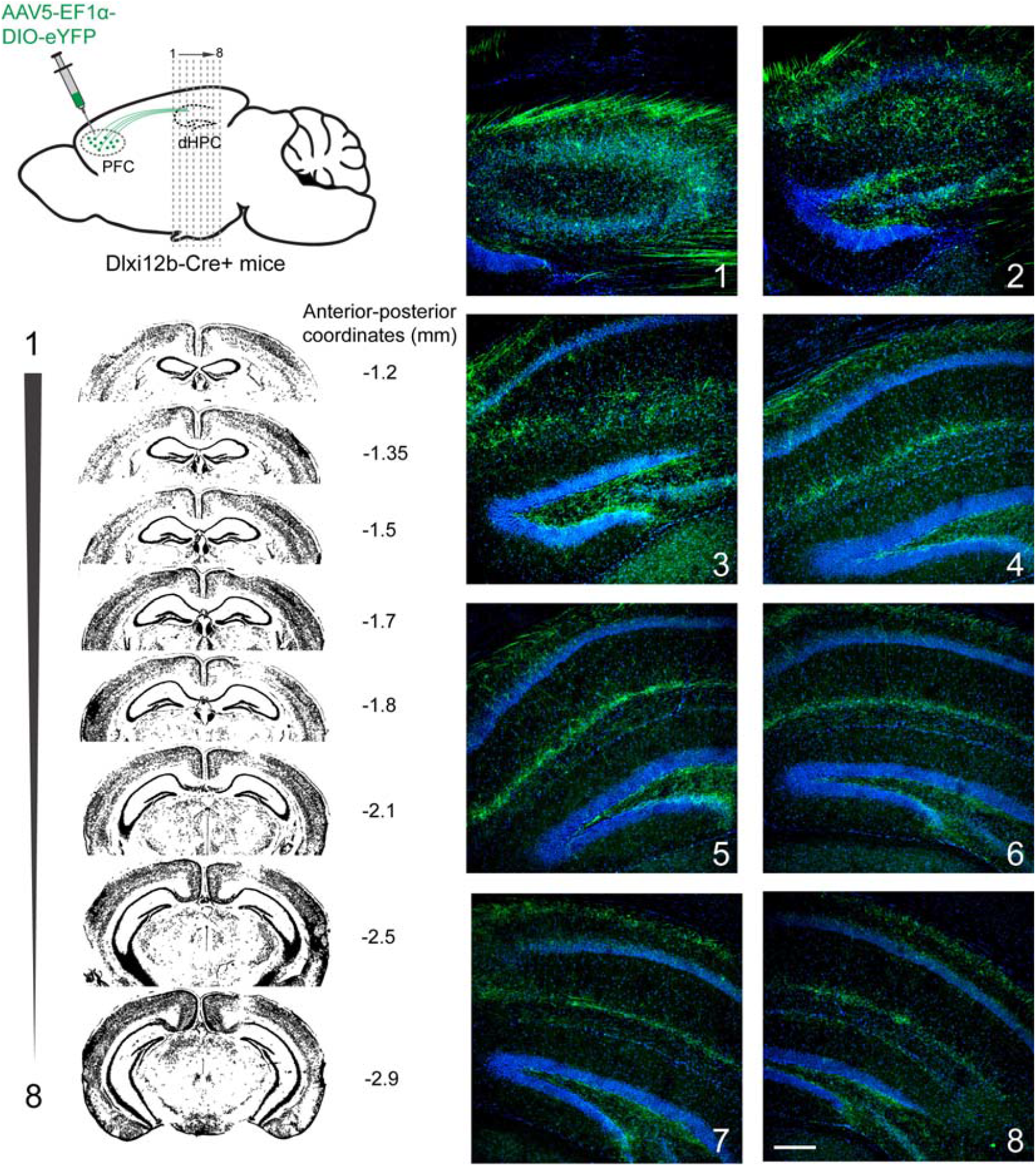
Anterograde tracing of HPC projecting PFC LRG neurons. Left: Schematic illustrating the anterograde tracing strategy. Microinjection of Cre-dependent eYFP virus into the PFC of *Dlxi12b-Cre*+ mice. Coronal slices of the hippocampus were obtained at increasing anterior-posterior (AP) distance from Bregma. Right: DAPI stained hippocampal sections showing eYFP+ PFC LRG axon terminals (green). Numbers on the right indicate the anterior-posterior (AP) coordinates w.r.t. bregma. Scale bar, 200 µm.

**Figure S2, related to Figure 2:**
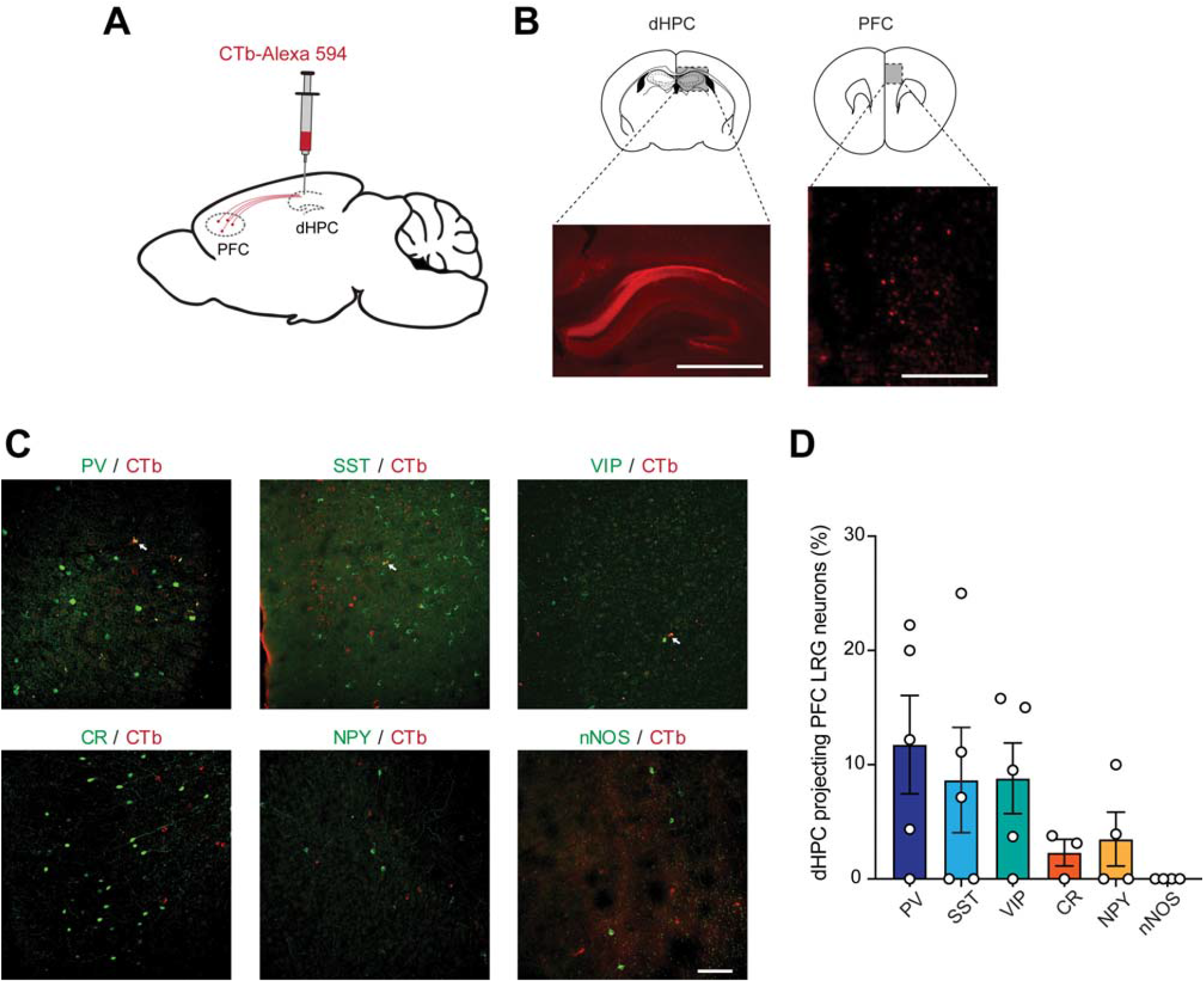
Inhibitory neuron marker expression in dHPC projecting PFC LRG neurons. **(A)** Schematic illustrating the experimental design: Retrograde tracer, Alexa 594 conjugated Cholera Toxin (CTb), was injected into dHPC. **(B)** Representative images showing Alexa 594-CTb injection site in dHPC section (left) and CTb+ neurons in PFC (right). Note: there is a high density of labeled axons in the corpus callosum. Scale bar, 1mm (left) and 250µm (right). **(C)** Example images showing co-labeling of inhibitory neuron markers (green) in CTb+ dHPC-projecting PFC LRG neurons. Parvalbumin (PV), Somatostatin (SST), Vasoactive intestinal polypeptide (VIP), Calretinin (CR), Neuropeptide Y (NPY), neuronal nitric oxide synthase (nNOS). Scale bar, 100 µm. **(D)** Percentage CTb+ dHPC projecting PFC LRG neurons co-expressing various inhibitory neuron markers (mean ± SEM). Empty circles represent values from individual sections.

**Figure S3, related to Figure 3:**
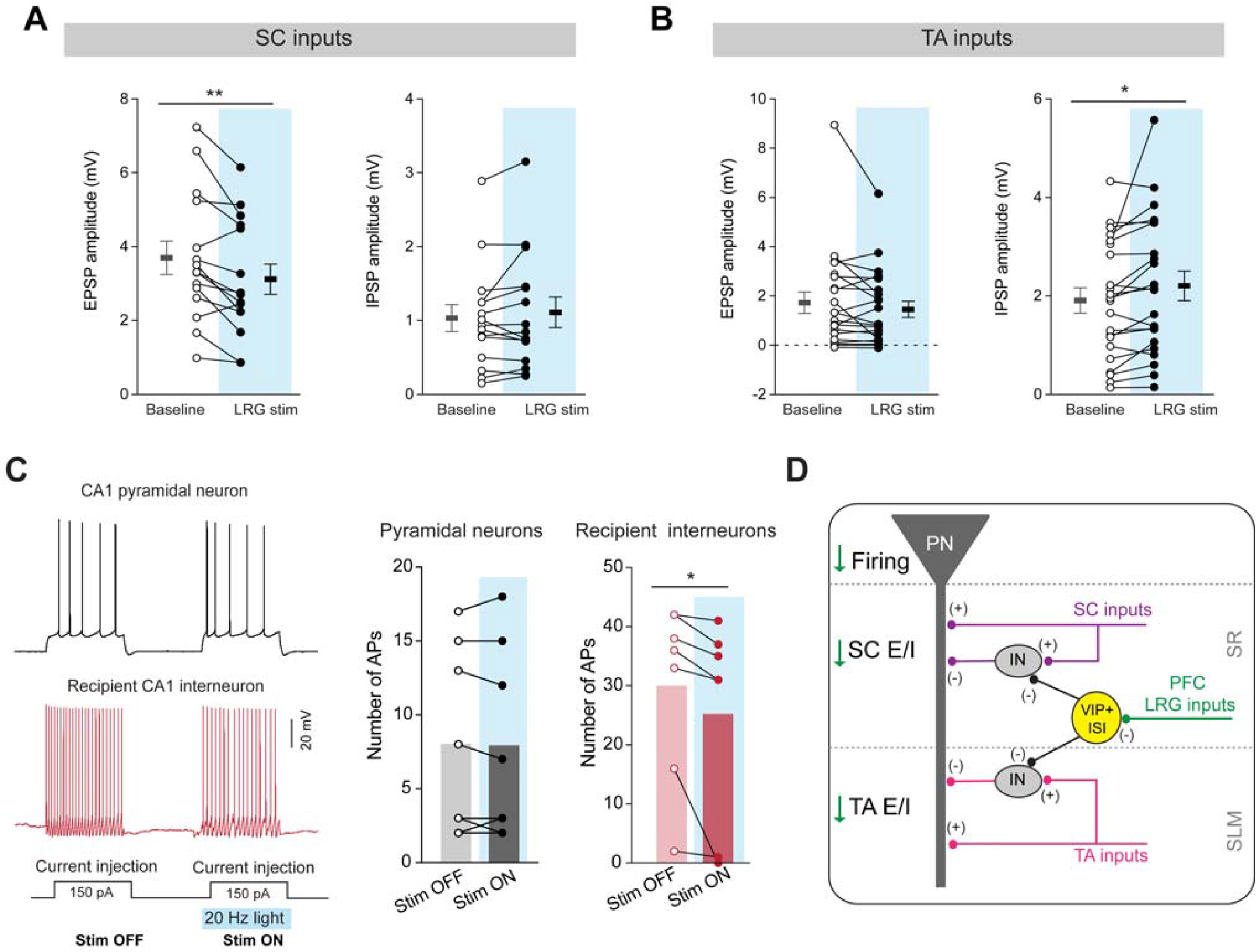
Dorsal HPC projecting PFC LRG neurons increase afferent input mediated feedforward inhibition in the CA1 microcircuit. **(A)** Amplitudes of SC (Schaffer collateral) pathway mediated EPSPs (left) and IPSPs (right) recorded in CA1 pyramidal neurons with (LRG stim) and without (Baseline) concomitant optogenetic stimulation of PFC LRG projections. Open and filled circles represent individual neurons (n = 15), horizontal lines represent averages (± SEM). Paired two-way t-test, ** p < 0.01. **(B)** Same as **A** for TA (temporoammonic) input pathway mediated EPSPs and IPSPs (n = 22). Paired two-way t-test, * p < 0.05. **(C)** Left: example voltage traces showing firing in response to depolarizing current injections with (Stim ON) and without (Stim OFF) concomitant optogenetic stimulation of PFC–dHPC LRG projections (470 nm, 5 ms pulses at 20 Hz; blue bar). Black trace is an example voltage response in CA1 pyramidal neuron. Red trace shows example voltage response in recipient CA1 interneuron. Right: firing output in the absence of optogenetic stimulation (Stim OFF) and during concomitant optogenetic stimulation (Stim ON) in CA1 pyramidal neurons (n = 10) and recipient CA1 interneurons (n = 7). Open and filled circles represent individual neurons, bars represent the average values. Two-way paired t-test; * p < 0.05. **(E)** Schematic illustrating the effects of PFC–dHPC LRG projection activation on microcircuit computation in dorsal CA1. PFC LRG projections inhibit VIP+ disinhibitory interneurons, and thereby increase SC and TA pathway mediated feedforward inhibition onto CA1 pyramidal neurons. The reduction in excitation-inhibition ratio at the afferent inputs reduces the afferent input mediated firing of CA1 pyramidal neurons.

**Figure S4, related to Figure 4:**
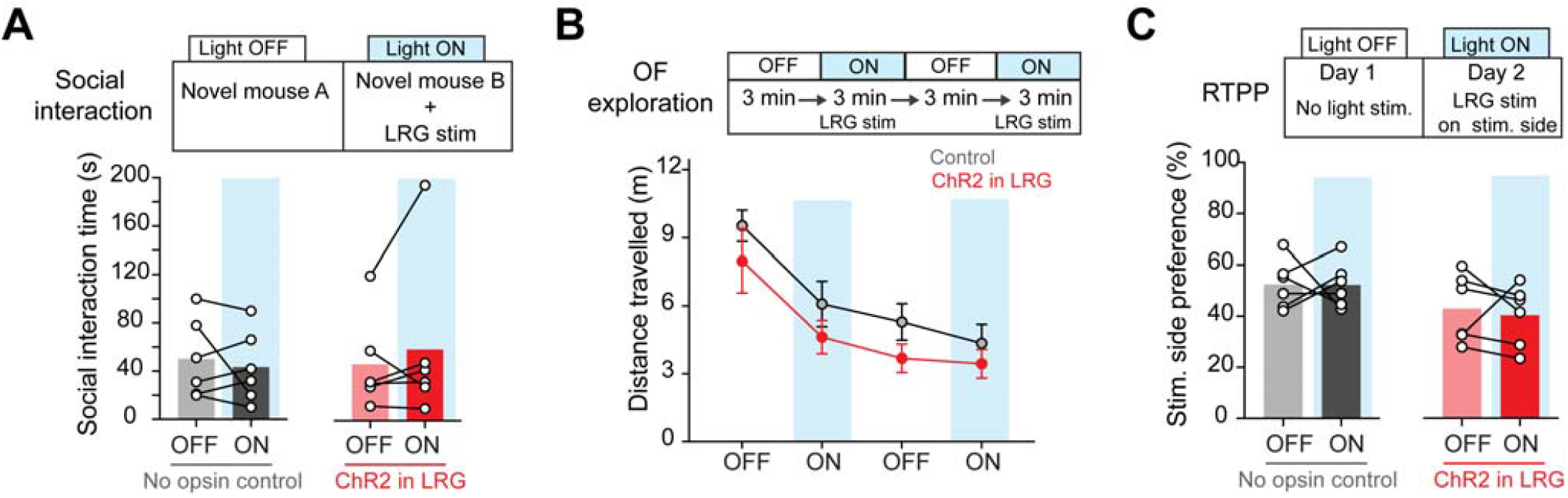
Optogenetic stimulation of PFC–dHPC LRG projections during social and exploratory behaviors. **(A)** Top: mice were tested for social interaction behavior with and without optogenetic stimulation. Bottom: Time spent interacting with a novel juvenile mouse is plotted for control and ChR2-expressing mice during light OFF and light ON periods. Open circles represent data from individual mice and bars represent the average values. **(B)** Top: During open-field (OF) exploration, PFC–dHPC LRG projections were optogenetically stimulated during the second and fourth 3-min epochs of the testing session. Bottom: Average (± SEM) distance travelled during OF exploration during light ON and light OFF epochs is plotted for control and ChR2-expressing mice. **(C)** Top: experimental design for real-time place preference (RTPP) test. Bottom: Preference for the stimulated chamber during light OFF and light ON conditions (expressed as a % of the total time) is plotted for control and ChR2-expressing mice. Open circles represent data from individual mice and bars represent the average values.

**Figure S5, related to Figure 5:**
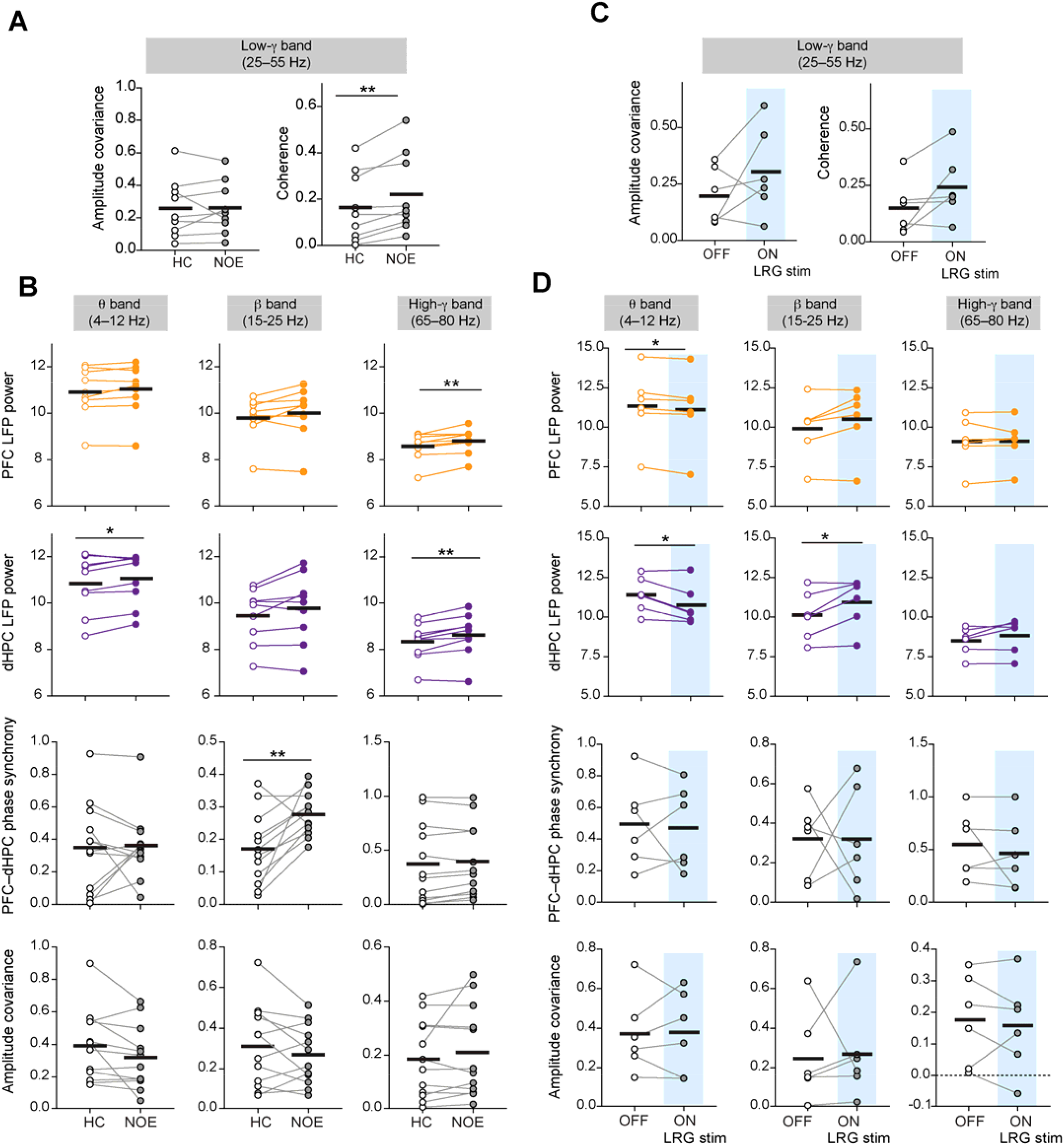
Effect of optogenetic stimulation of PFC–dHPC LRG projections and NOE on oscillatory activity in dHPC and PFC. **(A)** Low-gamma (low-γ) amplitude covariance and coherence between PFC and dHPC LFPs recorded during home cage exploration (HC) and novel object exploration (NOE) epochs. Open and filled circles represent data from individual mice (n = 9) and solid black lines represent average values. Two-way paired t-test, ** p < 0.01. **(B)** PFC and dHPC LFP log power, PFC-dHPC phase synchrony, and PFC-dHPC amplitude covariance for the theta (θ), beta (β) and high gamma (high-γ) frequency bands during HC and NOE epochs are plotted. Open and filled circles represent data from individual mice (n = 9) and solid black lines represent average values. Two-way paired t-test; * p < 0.05, **p < 0.01. **(C)** Low-γ amplitude covariance and coherence between PFC and dHPC LFPs recorded during Light OFF and Light ON epochs (optogenetic stimulation; 473 nm, 5 ms pulses at 20 Hz; blue bars). Open and filled circles represent data from individual mice (n = 6) and solid black lines represent average values. **(D)** PFC and dHPC LFP log power, PFC-dHPC phase synchrony, and PFC-dHPC amplitude covariance for the theta (θ), beta (β) and high-gamma (high-γ) frequency bands during Light OFF and Light ON epochs are plotted. Open and filled circles represent data from individual mice (n = 6) and solid black lines represent average values. Two-way paired t-test; * p < 0.05.

**Figure S6, related to Figure 6:**
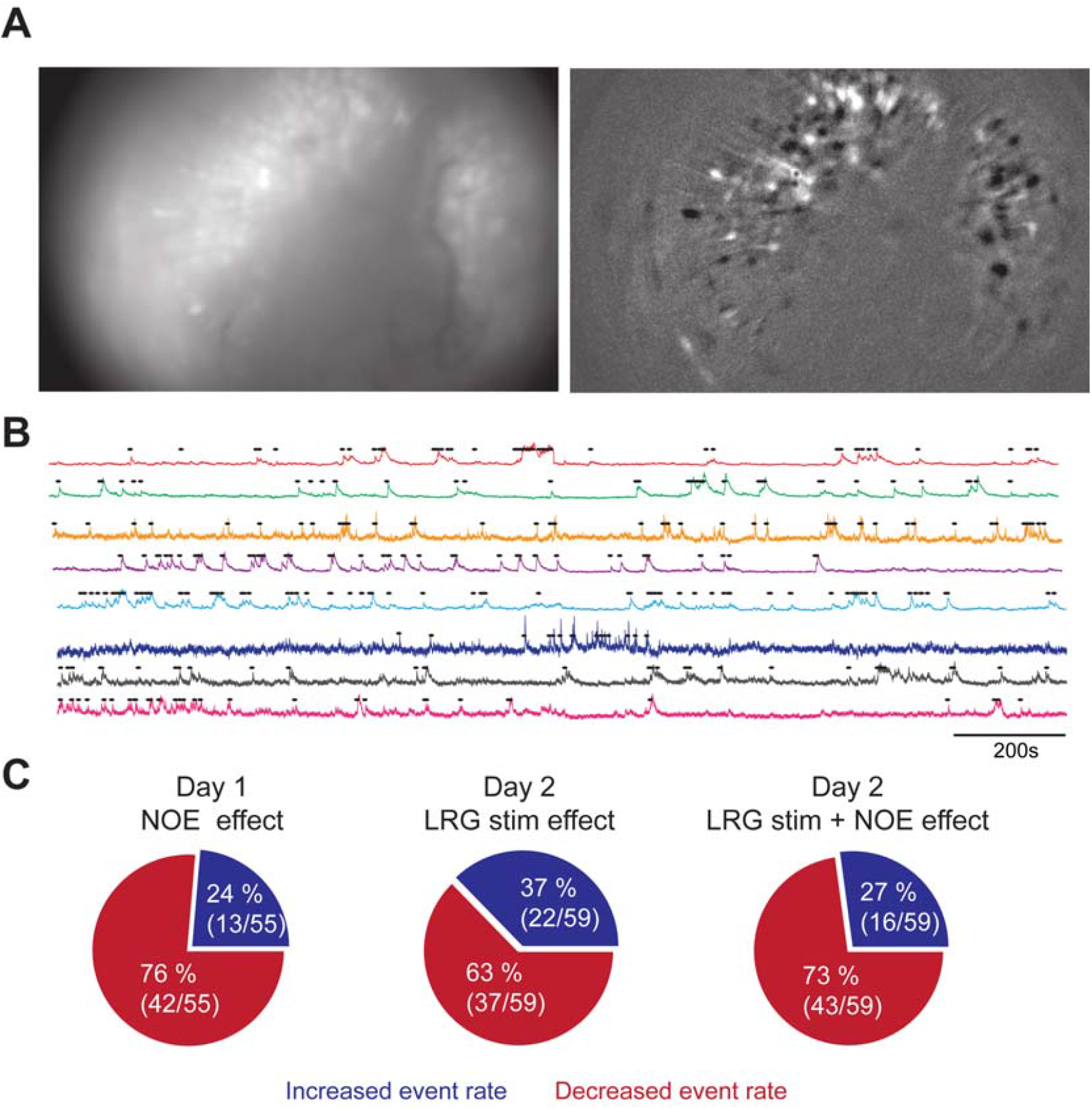
*In vivo* Ca^2+^ activity of dorsal CA1 neurons during optogenetic stimulation of PFC LRG projections. **(A)** Left: Representative raw epifluorescence image showing jGCaMP7f expression in the dorsal CA1 imaged using a miniaturized microscope in a freely behaving mouse. Right: Transformed image showing relative change in fluorescence (dF/F). **(B)** Representative Ca^2+^ transients (colored lines) recorded from dorsal CA1 neurons. Black dots above the traces represent the detected active Ca^2+^ events. **(C)** Pie charts showing the fraction of CA1 neurons that decrease or increase activity during NOE on day 1 testing, during LRG stimulation on day 2 testing, or during NOE in the presence of LRG stimulation on day 2 testing.

## SUPPLEMENTAL INFORMATION: TABLES

**Table S1,related to Figure 1:**
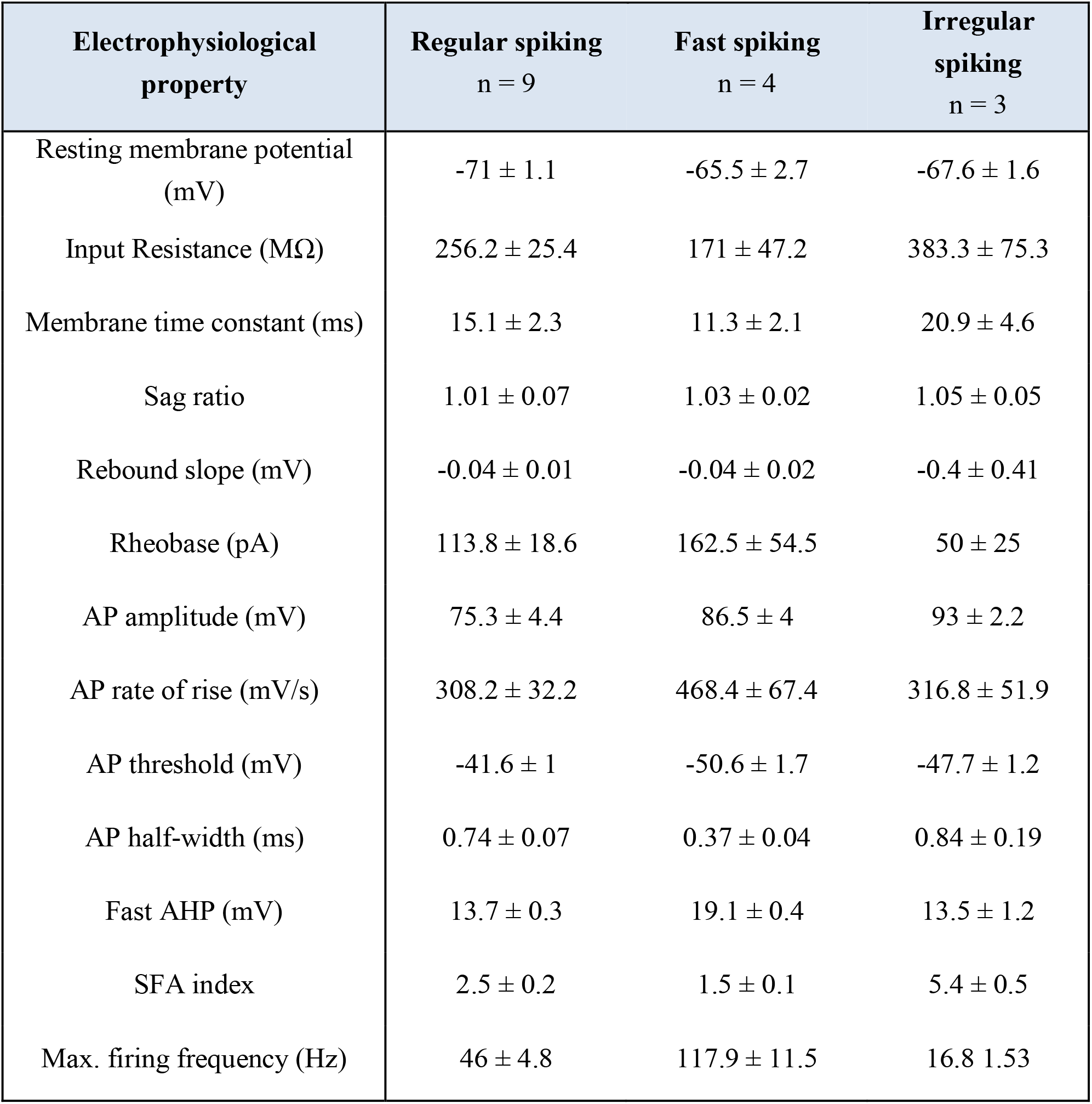
Electrophysiological heterogeneity of dHPC projecting PFC LRG neurons. Mean (± SEM) values for resting membrane properties, action-potential (AP) properties, and firing properties of dHPC projecting PFC LRG neurons classified as regular spiking, fast spiking, and irregular spiking (Fig. 1H). AHP, afterhyperpolarization; SFA, spike frequency accommodation.

**Table S2,related to Figure 2:**
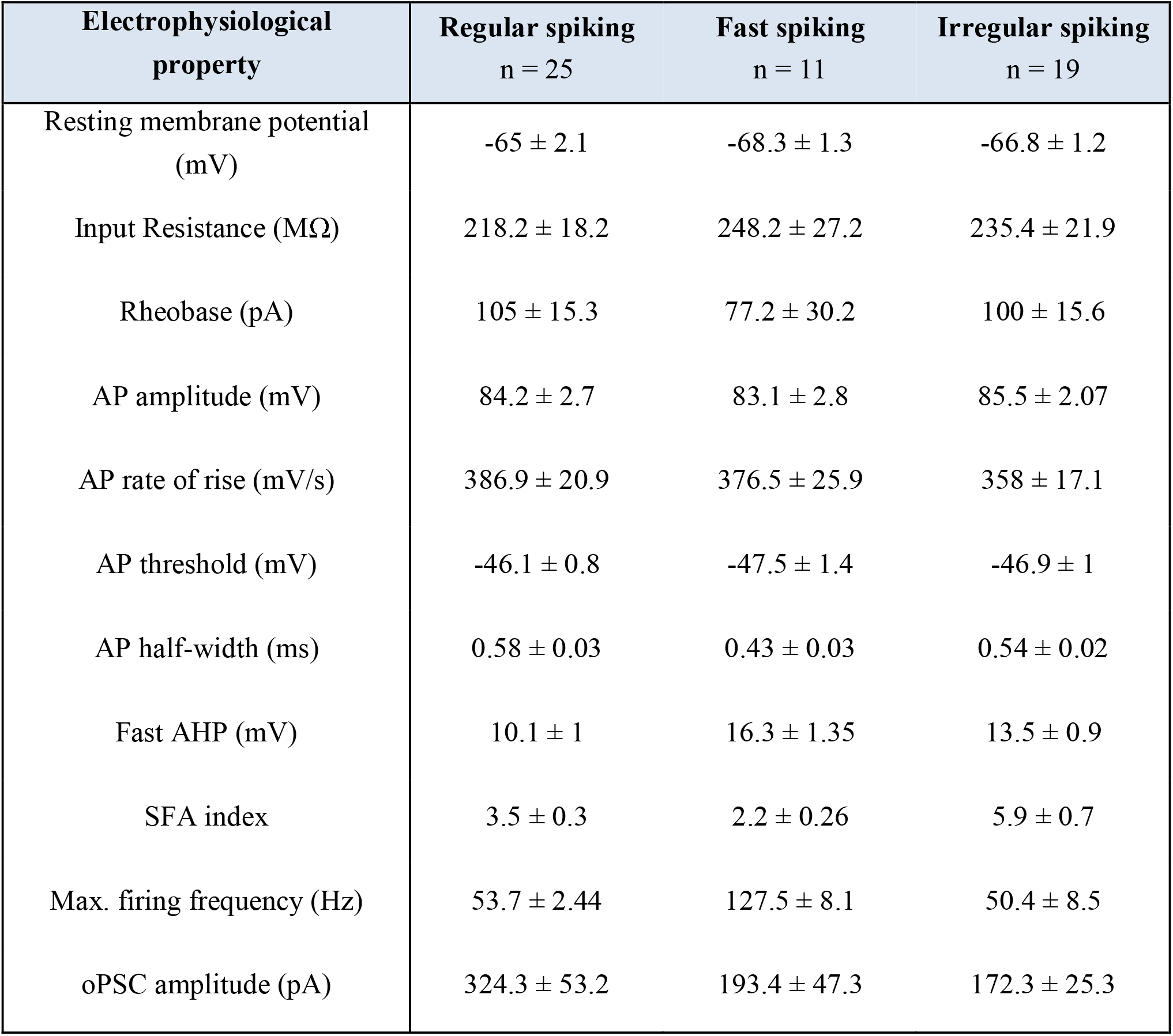
Electrophysiological properties of dHPC interneurons receiving direct PFC LRG inputs. Mean (± SEM) values for resting membrane properties, action-potential (AP) properties, firing properties, and optogenetically evoked postsynaptic current (oPSC) amplitudes recorded from recipient CA1 interneurons classified as regular spiking, fast spiking, and irregular spiking (Fig. 2B). AHP, afterhyperpolarization; SFA, spike frequency accommodation.

| Electrophysiological property | Regular spiking<br>n = 25 | Fast spiking<br>n = 11 | Irregular spiking<br>n = 19 |
| --- | --- | --- | --- |
| Resting membrane potential (mV) | -65 ± 2.1 | -68.3 ± 1.3 | -66.8 ± 1.2 |
| Input Resistance (MΩ) | 218.2 ± 18.2 | 248.2 ± 27.2 | 235.4 ± 21.9 |
| Rheobase (pA) | 105 ± 15.3 | 77.2 ± 30.2 | 100 ± 15.6 |
| AP amplitude (mV) | 84.2 ± 2.7 | 83.1 ± 2.8 | 85.5 ± 2.07 |
| AP rate of rise (mV/s) | 386.9 ± 20.9 | 376.5 ± 25.9 | 358 ± 17.1 |
| AP threshold (mV) | -46.1 ± 0.8 | -47.5 ± 1.4 | -46.9 ± 1 |
| AP half-width (ms) | 0.58 ± 0.03 | 0.43 ± 0.03 | 0.54 ± 0.02 |
| Fast AHP (mV) | 10.1 ± 1 | 16.3 ± 1.35 | 13.5 ± 0.9 |
| SFA index | 3.5 ± 0.3 | 2.2 ± 0.26 | 5.9 ± 0.7 |
| Max. firing frequency (Hz) | 53.7 ± 2.44 | 127.5 ± 8.1 | 50.4 ± 8.5 |
| oPSC amplitude (pA) | 324.3 ± 53.2 | 193.4 ± 47.3 | 172.3 ± 25.3 |

## REFERENCES

Acsády, L., Arabadzisz, D., and Freund, T.F. (1996a). Correlated morphological and neurochemical features identify different subsets of vasoactive intestinal polypeptide-immunoreactive interneurons in rat hippocampus. Neuroscience 73, 299–315.

Acsády, L., Görcs, T.J., and Freund, T.F. (1996b). Different populations of vasoactive intestinal polypeptide-immunoreactive interneurons are specialized to control pyramidal cells or interneurons in the hippocampus. Neuroscience 73, 317–334.

Ang, C.W., Carlson, G.C., and Coulter, D.A. (2005). Hippocampal CA1 Circuitry Dynamically Gates Direct Cortical Inputs Preferentially at Theta Frequencies. J Neurosci 25, 9567–9580.

Basu, J., Zaremba, J.D., Cheung, S.K., Hitti, F.L., Zemelman, B.V., Losonczy, A., and Siegelbaum, S.A. (2016). Gating of hippocampal activity, plasticity, and memory by entorhinal cortex long-range inhibition. Science 351, aaa5694–aaa5694.

Bähner, F., Demanuele, C., Schweiger, J., Gerchen, M.F., Zamoscik, V., Ueltzhöffer, K., Hahn, T., Meyer, P., Flor, H., Durstewitz, D., et al. (2015). Hippocampal-dorsolateral prefrontal coupling as a species-conserved cognitive mechanism: a human translational imaging study. Neuropsychopharmacology 40, 1674–1681.

Bittner, K.C., Grienberger, C., Vaidya, S.P., Milstein, A.D., Macklin, J.J., Suh, J., Tonegawa, S., and Magee, J.C. (2015). Conjunctive input processing drives feature selectivity in hippocampal CA1 neurons. Nat Neurosci 18, 1133–1142.

Brincat, S.L., and Miller, E.K. (2015). Frequency-specific hippocampal-prefrontal interactions during associative learning. Nat Neurosci 18, 576–581.

Chamberland, S., and Topolnik, L. (2012). Inhibitory control of hippocampal inhibitory neurons. Front. Neurosci. 6, 165.

Chattopadhyaya, B., and Cristo, G.D. (2012). GABAergic circuit dysfunctions in neurodevelopmental disorders. Front Psychiatry 3, 51–51.

Christenson Wick, Z., Tetzlaff, M.R., and Krook-Magnuson, E. (2019). Novel long-range inhibitory nNOS-expressing hippocampal cells. eLife 8, e46816.

Churchwell, J.C., Morris, A.M., Musso, N.D., and Kesner, R.P. (2010). Prefrontal and hippocampal contributions to encoding and retrieval of spatial memory. Neurobiology of Learning and Memory 93, 415–421.

Colgin, L.L. (2011). Oscillations and hippocampal–prefrontal synchrony. Current Opinion in Neurobiology 21, 467–474.

Corder, G., Ahanonu, B., Grewe, B.F., Wang, D., Schnitzer, M.J., and Scherrer, G. (2019). An amygdalar neural ensemble that encodes the unpleasantness of pain. Science 363, 276.

Csicsvari, J., Jamieson, B., Wise, K.D., and Buzsáki, G. (2003). Mechanisms of Gamma Oscillations in the Hippocampus of the Behaving Rat. Neuron 37, 311–322.

Cunha-Reis, D., and Caulino-Rocha, A. (2020). VIP Modulation of Hippocampal Synaptic Plasticity: A Role for VIP Receptors as Therapeutic Targets in Cognitive Decline and Mesial Temporal Lobe Epilepsy. Front Cell Neurosci 14, 153–153.

Cunniff, M.M., Markenscoff-Papadimitriou, E., Ostrowski, J., Rubenstein, J.L., and Sohal, V.S. (2020). Altered hippocampal-prefrontal communication during anxiety-related avoidance in mice deficient for the autism-associated gene Pogz. eLife 9, e54835.

Dana, H., Sun, Y., Mohar, B., Hulse, B.K., Kerlin, A.M., Hasseman, J.P., Tsegaye, G., Tsang, A., Wong, A., Patel, R., et al. (2019). High-performance calcium sensors for imaging activity in neuronal populations and microcompartments. Nat Meth 16, 649–657.

del Rio, D., Beucher, B., Lavigne, M., Wehbi, A., Gonzalez Dopeso-Reyes, I., Saggio, I., and Kremer, E.J. (2019). CAV-2 Vector Development and Gene Transfer in the Central and Peripheral Nervous Systems. Front. Mol. Neurosci. 12, 1202.

DeVito, L.M., and Eichenbaum, H. (2010). Distinct contributions of the hippocampus and medial prefrontal cortex to the “what-where-when” components of episodic-like memory in mice. Behav Brain Res 215, 318–325.

Eichenbaum, H. (2017). Prefrontal–hippocampal interactions in episodic memory. Nat Rev Neurosci 18, 547–558.

Fanselow, M.S., and Dong, H.-W. (2010). Are the Dorsal and Ventral Hippocampus Functionally Distinct Structures? Neuron 65, 7–19.

Floresco, S.B., Seamans, J.K., and Phillips, A.G. (1997). Selective Roles for Hippocampal, Prefrontal Cortical, and Ventral Striatal Circuits in Radial-Arm Maze Tasks With or Without a Delay. J Neurosci 17, 1880.

Francavilla, R., Villette, V., Luo, X., Chamberland, S., Muñoz-Pino, E., Camiré, O., Wagner, K., Kis, V., Somogyi, P., and Topolnik, L. (2018). Connectivity and network state-dependent recruitment of long-range VIP-GABAergic neurons in the mouse hippocampus. Nature Communications 9, 5043.

Frost, N.A., Haggart, A., and Sohal, V.S. (2020). Dynamic correlations help prefrontal ensembles transmit information about social behavior. bioRxiv 2020.08.05.238741.

Gazzaley, A., and D’Esposito, M. (2007). Unifying prefrontal cortex function: Executive control, neural networks, and top-down modulation. In The Human Frontal Lobes: Functions and Disorders, 2nd Ed, (New York, NY, US: The Guilford Press), pp. 187–206.

Gilbert, S.J., Bird, G., Brindley, R., Frith, C.D., and Burgess, P.W. (2008). Atypical recruitment of medial prefrontal cortex in autism spectrum disorders: An fMRI study of two executive function tasks. Neuropsychologia 46, 2281–2291.

Godsil, B.P., Kiss, J.P., Spedding, M., and Jay, T.M. (2013). The hippocampal-prefrontal pathway: the weak link in psychiatric disorders? European Neuropsychopharmacology 23, 1165–1181.

Grienberger, C., Milstein, A.D., Bittner, K.C., Romani, S., and Magee, J.C. (2017). Inhibitory suppression of heterogeneously tuned excitation enhances spatial coding in CA1 place cells. Nat Neurosci 20, 417–426.

Guise, K.G., and Shapiro, M.L. (2017). Medial Prefrontal Cortex Reduces Memory Interference by Modifying Hippocampal Encoding. Neuron 94, 183–192.e188.

Hallock, H.L., Wang, A., and Griffin, A.L. (2016). Ventral Midline Thalamus Is Critical for Hippocampal-Prefrontal Synchrony and Spatial Working Memory. J. Neurosci 36, 8372–8389.

Hare, B.D., and Duman, R.S. (2020). Prefrontal cortex circuits in depression and anxiety: contribution of discrete neuronal populations and target regions. Molecular Psychiatry 2014 20:7 25, 2742–2758.

Hnasko, T.S., Perez, F.A., Scouras, A.D., Stoll, E.A., Gale, S.D., Luquet, S., Phillips, P.E.M., Kremer, E.J., and Palmiter, R.D. (2006). Cre recombinase-mediated restoration of nigrostriatal dopamine in dopamine-deficient mice reverses hypophagia and bradykinesia. Proc Natl Acad Sci USA 103, 8858.

Hoover, W.B., and Vertes, R.P. (2007). Anatomical analysis of afferent projections to the medial prefrontal cortex in the rat. Brain Struct Funct 212, 149–179.

Hoover, W.B., and Vertes, R.P. (2012). Collateral projections from nucleus reuniens of thalamus to hippocampus and medial prefrontal cortex in the rat: a single and double retrograde fluorescent labeling study. Brain Struct Funct 217, 191–209.

Jay, T.R.S.M., and Witter, M.P. (1991). Distribution of hippocampal CA1 and subicular efferents in the prefrontal cortex of the rat studied by means of anterograde transport ofPhaseolus vulgaris-leucoagglutinin. J. Comp. Neurol. 313, 574–586.

Jin, J., and Maren, S. (2015). Prefrontal-Hippocampal Interactions in Memory and Emotion. Front. Syst. Neurosci. 9, 1082.

Jinno, S., Klausberger, T., Marton, L.F., Dalezios, Y., Roberts, J.D.B., Fuentealba, P., Bushong, E.A., Henze, D., Buzsaki, G., and Somogyi, P. (2007). Neuronal Diversity in GABAergic Long-Range Projections from the Hippocampus. J. Neurosci 27, 8790–8804.

Jones, M.W., and Wilson, M.A. (2005). Theta Rhythms Coordinate Hippocampal–Prefrontal Interactions in a Spatial Memory Task. PLoS Biol 3, e402.

Klapoetke, N.C., Murata, Y., Kim, S.S., Pulver, S.R., Birdsey-Benson, A., Cho, Y.K., Morimoto, T.K., Chuong, A.S., Carpenter, E.J., Tian, Z., et al. (2014). Independent optical excitation of distinct neural populations. Nat Meth 11, 338–346.

Kupferschmidt, D.A., and Gordon, J.A. (2018). The dynamics of disordered dialogue: Prefrontal, hippocampal and thalamic miscommunication underlying working memory deficits in schizophrenia. Brain and Neuroscience Advances 2, 239821281877182.

Kyd, R.J., and Bilkey, D.K. (2003). Prefrontal Cortex Lesions Modify the Spatial Properties of Hippocampal Place Cells. Cerebral Cortex 13, 444–451.

Lee, A.T., Cunniff, M.M., See, J.Z., Wilke, S.A., Luongo, F.J., Ellwood, I.T., Ponnavolu, S., and Sohal, V.S. (2019). VIP Interneurons Contribute to Avoidance Behavior by Regulating Information Flow across Hippocampal-Prefrontal Networks. Neuron 102, 1223–1234.e1224.

Lee, A.T., Vogt, D., Rubenstein, J.L., and Sohal, V.S. (2014). A Class of GABAergic Neurons in the Prefrontal Cortex Sends Long-Range Projections to the Nucleus Accumbens and Elicits Acute Avoidance Behavior. J Neurosci 34, 11519–11525.

Li, M., Long, C., and Yang, L. (2015). Hippocampal-Prefrontal Circuit and Disrupted Functional Connectivity in Psychiatric and Neurodegenerative Disorders. BioMed Research International 2015, 810548–810548.

Malik, R., and Johnston, D. (2017). Dendritic GIRK Channels Gate the Integration Window, Plateau Potentials, and Induction of Synaptic Plasticity in Dorsal But Not Ventral CA1 Neurons. J Neurosci 37, 3940.

Malik, R., Dougherty, K.A., Parikh, K., Byrne, C., and Johnston, D. (2015). Mapping the electrophysiological and morphological properties of CA1 pyramidal neurons along the longitudinal hippocampal axis. Hippocampus 26, 341–361.

Malik, R., Pai, E.L.-L., Rubin, A.N., Stafford, A.M., Angara, K., Minasi, P., Rubenstein, J.L., Sohal, V.S., and Vogt, D. (2019). Tsc1 represses parvalbumin expression and fast-spiking properties in somatostatin lineage cortical interneurons. Nature Communications 10, 4994.

Marin, O. (2012). Interneuron dysfunction in psychiatric disorders. Nat Rev Neurosci 13, 107–120.

Mathis, A., Mamidanna, P., Cury, K.M., Abe, T., Murthy, V.N., Mathis, M.W., and Bethge, M. (2018). DeepLabCut: markerless pose estimation of user-defined body parts with deep learning. Nat Neurosci 21, 1281–1289.

McKenzie, S. (2018). Inhibition shapes the organization of hippocampal representations. Hippocampus 28, 659–671.

Melzer, S., and Monyer, H. (2020). Diversity and function of corticopetal and corticofugal GABAergic projection neurons. Nat Rev Neurosci 21, 499–515.

Melzer, S., Michael, M., Caputi, A., Eliava, M., Fuchs, E.C., Whittington, M.A., and Monyer, H. (2012). Long-range-projecting GABAergic neurons modulate inhibition in hippocampus and entorhinal cortex. Science 335, 1506–1510.

Miller, E.K., and Cohen, J.D. (2001). An integrative theory of prefrontal cortex function. Annu. Rev. Neurosci. 24, 167–202.

Miller, E.K. (2000). The prefontral cortex and cognitive control. Nat Rev Neurosci 1, 59–65.

Milstein, A.D., Bloss, E.B., Apostolides, P.F., Vaidya, S.P., Dilly, G.A., Zemelman, B.V., and Magee, J.C. (2015). Inhibitory Gating of Input Comparison in the CA1 Microcircuit. Neuron 87, 1274–1289.

Moser, E.I., Kropff, E., and Moser, M.-B. (2008). Place Cells, Grid Cells, and the Brain’s Spatial Representation System. Annu. Rev. Neurosci. 31, 69–89.

Mukamel, E.A., Nimmerjahn, A., and Schnitzer, M.J. (2009). Automated analysis of cellular signals from large-scale calcium imaging data. Neuron 63, 747–760.

O’Keefe, J. (1976). Place units in the hippocampus of the freely moving rat. Experimental Neurology 51, 78–109.

O’Neill, P.K., Gordon, J.A., and Sigurdsson, T. (2013). Theta Oscillations in the Medial Prefrontal Cortex Are Modulated by Spatial Working Memory and Synchronize with the Hippocampus through Its Ventral Subregion. J. Neurosci 33, 14211–14224.

Orellana, G., and Slachevsky, A. (2013). Executive Functioning in Schizophrenia. Front Psychiatry 4, 35.

Paterno, R., Casalia, M., and Baraban, S.C. (2020). Interneuron deficits in neurodevelopmental disorders: Implications for disease pathology and interneuron-based therapies. European Journal of Paediatric Neurology 24, 81–88.

Place, R., Farovik, A., Brockmann, M., and Eichenbaum, H. (2016). Bidirectional prefrontal-hippocampal interactions support context-guided memory. Nat Neurosci 19, 992–994.

Potter, G.B., Petryniak, M.A., Shevchenko, E., McKinsey, G.L., Ekker, M., and Rubenstein, J.L.R. (2009). Generation of Cre-transgenic mice using Dlx1/Dlx2 enhancers and their characterization in GABAergic interneurons. 40, 167–186.

Preston, A.R., and Eichenbaum, H. (2013). Interplay of hippocampus and prefrontal cortex in memory. Current Biology 23, R764–R773.

Rajasethupathy, P., Sankaran, S., Marshel, J.H., Kim, C.K., Ferenczi, E., Lee, S.Y., Berndt, A., Ramakrishnan, C., Jaffe, A., Lo, M., et al. (2015). Projections from neocortex mediate top-down control of memory retrieval. Nature 526, 653–659.

Shin, J.D., and Jadhav, S.P. (2016). Multiple modes of hippocampal–prefrontal interactions in memory-guided behavior. Systems Neuroscience 40, 161–169.

Sigurdsson, T., and Duvarci, S. (2016). Hippocampal-Prefrontal Interactions in Cognition, Behavior and Psychiatric Disease. Front. Syst. Neurosci. 9, 190–190.

Sigurdsson, T., Stark, K.L., Karayiorgou, M., Gogos, J.A., and Gordon, J.A. (2010). Impaired hippocampal–prefrontal synchrony in a genetic mouse model of schizophrenia. Nature 464, 763–767.

Spellman, T., Rigotti, M., Ahmari, S.E., Fusi, S., Gogos, J.A., and Gordon, J.A. (2015). Hippocampal-prefrontal input supports spatial encoding in working memory. Nature 522, 309–314.

Stamatakis, A.M., Schachter, M.J., Gulati, S., Zitelli, K.T., Malanowski, S., Tajik, A., Fritz, C., Trulson, M., and Otte, S.L. (2018). Simultaneous Optogenetics and Cellular Resolution Calcium Imaging During Active Behavior Using a Miniaturized Microscope. Front. Neurosci. 12, 496–496.

Tamamaki, N., and Tomioka, R. (2010). Long-Range GABAergic Connections Distributed throughout the Neocortex and their Possible Function. Front. Neurosci. 4, 202.

Trimper, J.B., Galloway, C.R., Jones, A.C., Mandi, K., and Manns, J.R. (2017). Gamma Oscillations in Rat Hippocampal Subregions Dentate Gyrus, CA3, CA1, and Subiculum Underlie Associative Memory Encoding. CellReports 21, 2419–2432.

Tukker, J.J., Fuentealba, P., Hartwich, K., Somogyi, P., and Klausberger, T. (2007). Cell Type-Specific Tuning of Hippocampal Interneuron Firing during Gamma Oscillations In Vivo. J Neurosci 27, 8184.

Turi, G.F., Li, W.-K., Chavlis, S., Pandi, I., O’Hare, J., Priestley, J.B., Grosmark, A.D., Liao, Z., Ladow, M., Zhang, J.F., et al. (2019). Vasoactive Intestinal Polypeptide-Expressing Interneurons in the Hippocampus Support Goal-Oriented Spatial Learning. Neuron 101, 1150–1165.e1158.

Vertes, R.P., Hoover, W.B., Szigeti-Buck, K., and Leranth, C. (2007). Nucleus reuniens of the midline thalamus: Link between the medial prefrontal cortex and the hippocampus. Brain Research Bulletin 71, 601–609.

Vinck, M., Oostenveld, R., van Wingerden, M., Battaglia, F., and Pennartz, C.M.A. (2011). An improved index of phase-synchronization for electrophysiological data in the presence of volume-conduction, noise and sample-size bias. Neuroimage 55, 1548–1565.

Wang, G.-W., and Cai, J.-X. (2006). Disconnection of the hippocampal–prefrontal cortical circuits impairs spatial working memory performance in rats. Behavioural Brain Research 175, 329–336.

Wilson, M.A., and McNaughton, B.L. (1993). Dynamics of the hippocampal ensemble code for space. Science 261, 1055.

Xu, W., and Südhof, T.C. (2013). A neural circuit for memory specificity and generalization. Science 339, 1290–1295.

Yoon, T., Okada, J., Jung, M.W., and Kim, J.J. (2008). Prefrontal cortex and hippocampus subserve different components of working memory in rats. Learn. Mem. 15, 97–105.

Yu, J.Y., and Frank, L.M. (2015). Hippocampal–cortical interaction in decision making. Neurobiology of Learning and Memory 117, 34–41.

